# Flow-cell based technology for massively parallel characterization of base-modified DNA aptamers

**DOI:** 10.1101/2020.04.25.060004

**Authors:** Diana Wu, Trevor Feagin, Peter Mage, Alexandra Rangel, Leighton Wan, Dehui Kong, Anping Li, John Coller, Michael Eisenstein, H. Tom Soh

## Abstract

Aptamers incorporating chemically modified bases can achieve superior affinity and specificity compared to natural aptamers, but their characterization remains a labor-intensive, low-throughput task. Here we describe the ‘non-natural aptamer array’ (N2A2) system, in which a minimally modified Illumina MiSeq instrument is used for the high-throughput generation and characterization of large libraries (∼10^6^) of base-modified DNA aptamer candidates on the basis of both target affinity and specificity. We first demonstrate the capability to screen multiple different base modifications to identify the optimal chemistry for high-affinity target binding. We next use N2A2 to generate aptamers that can maintain excellent specificity even in complex samples, with equally strong target affinity in both buffer and diluted human serum. Given that N2A2 requires only minor mechanical modifications to the MiSeq, we believe N2A2 offers a broadly accessible tool for generating high-quality affinity reagents for diverse applications.

## Introduction

Base-modified aptamers, in which natural DNA or RNA bases are replaced with non-natural, chemically-modified counterparts, can greatly enhance the affinity and specificity of aptamers for protein targets. For example, pioneering work by Gold and co-workers showed that natural aptamers can only target < 30% of the human proteome, whereas base-modified aptamers can target more than 80%.^1^ Other reports^2,3^ have shown that that the addition of non-natural functional groups at the 5-position of pyrimidines, such as a hydrophobic pentynyl^4^ or a cationic amine^5^, can result in aptamers that exhibit superior affinity or specificity to natural DNA aptamers. Base-modified aptamers can incorporate a wide range of chemical functional groups beyond those found in nature.^6^ This offers the exciting potential for generating affinity reagents for a broader spectrum of both protein and non-protein analytes that would otherwise be difficult or impossible to target.^7,8^ In the commercial arena, companies such as Somalogic has been successful in demonstrating the generation of many high-affinity ‘slow off-rate modified aptamers’ (SOMAmers) and shown the utility of these reagents for generating reproducible measurements of circulating proteins.^9,10^ Unfortunately, the sequences of these aptamers are not available to the research community and only a few base-modified aptamers have been reported in the literature by academic research groups ^2,3,11^.

Perhaps the most resource-intensive step of the base-modified aptamer discovery process is the binding characterization after the selection process. Historically, the affinity and specificity of individual aptamer candidates has been laboriously measured in a serial manner, but several groups have demonstrated higher-throughput platforms for the accelerated profiling of interactions between natural DNA or RNA and protein targets. For example, our group used a custom DNA microarray to characterize the affinity and specificity of ∼1,000 DNA aptamers that bind angiopoietin-2.^12^ The Burge lab demonstrated the use of flow-cells from a high-throughput sequencing (HTS) instrument, in which an Illumina Genome Analyzer (GAII) was used to sequence DNA clusters and measure protein binding on the same flow-cell^13^. This enabled them to thoroughly characterize elements influencing binding of the transcription factor Gcn4p to its consensus target DNA sequence. Jung *et al*. also employed a flow-cell-based screening approach to characterize the DNA sequence binding preference of CRISPR-Cas complexes.^14^ To enable a similar approach for RNA, the Lis group has developed a GAII-based strategy in which the instrument sequenced DNA within the flow-cell, transcribed each cluster into RNA *in situ*, used the terminator protein Tus to halt transcription, and then measured protein binding to the tethered RNA^15^. This allowed them to study the affinities and binding determinants of mutagenized pools of GFP and NELF E aptamers. The Greenleaf lab developed a similar HTS-based *in situ* transcription approach termed RNA-MaP,^16^ which they used to study the affinity of the MS2 coat protein for >10^7^ RNA targets. These various systems demonstrate the feasibility of re-purposing HTS instruments for parallel aptamer characterization, but there has been no equivalent demonstration of a strategy for characterizing base-modified aptamers in a high-throughput manner.

In this work, we describe a versatile platform that enables the efficient characterization of large numbers of base-modified DNA aptamers in a parallel fashion. Our ‘non-natural aptamer array’ (N2A2) system uses a modified benchtop Illumina MiSeq instrument to generate >10^6^ base-modified aptamer sequences *in situ* within the flow-cell, and subsequently measures the affinity and specificity of these sequences in parallel. N2A2 uses a generalizable copper-catalyzed azide-alkyne cycloaddition (CuAAC) ‘click chemistry’-based approach, initially described by Tolle *et al*.^17^, which enables efficient incorporation of virtually any chemical modification into our aptamers with commercially available polymerase enzymes. This simplifies the measurement and comparison of multiple modifications in the same pool of aptamers through consecutive sequencing experiments, each employing a different modification. Additionally, aptamer characterization can be performed directly in complex sample matrices, such that both specificity and affinity can be measured simultaneously. This enables efficient selection of modified aptamers that are optimally suited for a given sample.

As an initial demonstration, we used N2A2 to screen libraries containing different modifications in order to identify the best modification for generating aptamers that bind vascular endothelial growth factor (VEGF), a well-characterized protein that offers a useful benchmark for evaluating aptamer screening strategies. We compared the performance of libraries that were modified with either tyrosine or tryptophan, determined that tryptophan-modified libraries exhibit superior performance, and subsequently generated a tryptophan-modified aptamer with substantially better affinity than a previously published natural DNA VEGF aptamer.^18^ We also demonstrate the feasibility of screening for affinity and specificity simultaneously with N2A2. As an exemplar, we screened phenylalanine-modified aptamers that bind insulin in diluted serum. This small polypeptide is a challenging target – we are aware of only one insulin aptamer in the literature^19^ – and real world applications for insulin detection generally require robust affinity in serum samples. Using N2A2, we successfully obtained a base-modified aptamer that exhibits superior affinity to the previously reported aptamer in buffer and dramatically outperforms this molecule in serum. This work thus highlights the broad capabilities of N2A2 as a platform for characterizing based-modified aptamers.

## Results

### Overview of the N2A2 platform

N2A2 is designed to efficiently characterize the affinity and specificity of many base-modified aptamers from an enriched pool. The N2A2 is fabricated by converting the Illumina MiSeq instrument such that base-modified aptamer clusters can be synthesized and characterized directly on the sequencing flow-cell (**Fig. 1A**). The workflow entails three main steps: sequencing, conversion, and *in situ* binding measurements (**Fig. 1B)**. It begins with sequencing of a DNA library during read 1 of a paired-end sequencing run. During the paired-end turnaround and read 2,^20,21^ the DNA clusters are converted into base-modified aptamer clusters. This is achieved via a CuAAC-based click chemistry reaction^17,22^, which offers a generalizable approach for coupling azide-modified functional groups onto alkyne-modified nucleobases, which can be incorporated into aptamer sequences with a standard commercially-available polymerase. Finally, during the rest of read 2, the flow-cell is incubated with fluorescently-labeled protein to screen for affinity and specificity, with intensity information collected for each cluster. These sequencing and screening data are processed to generate a phenotype-genotype linked map of all the clusters. The initial libraries that we employed comprise a randomized region flanked by forward primer (FP) and reverse primer complementary (RPc) sequences.

**Figure 1.**
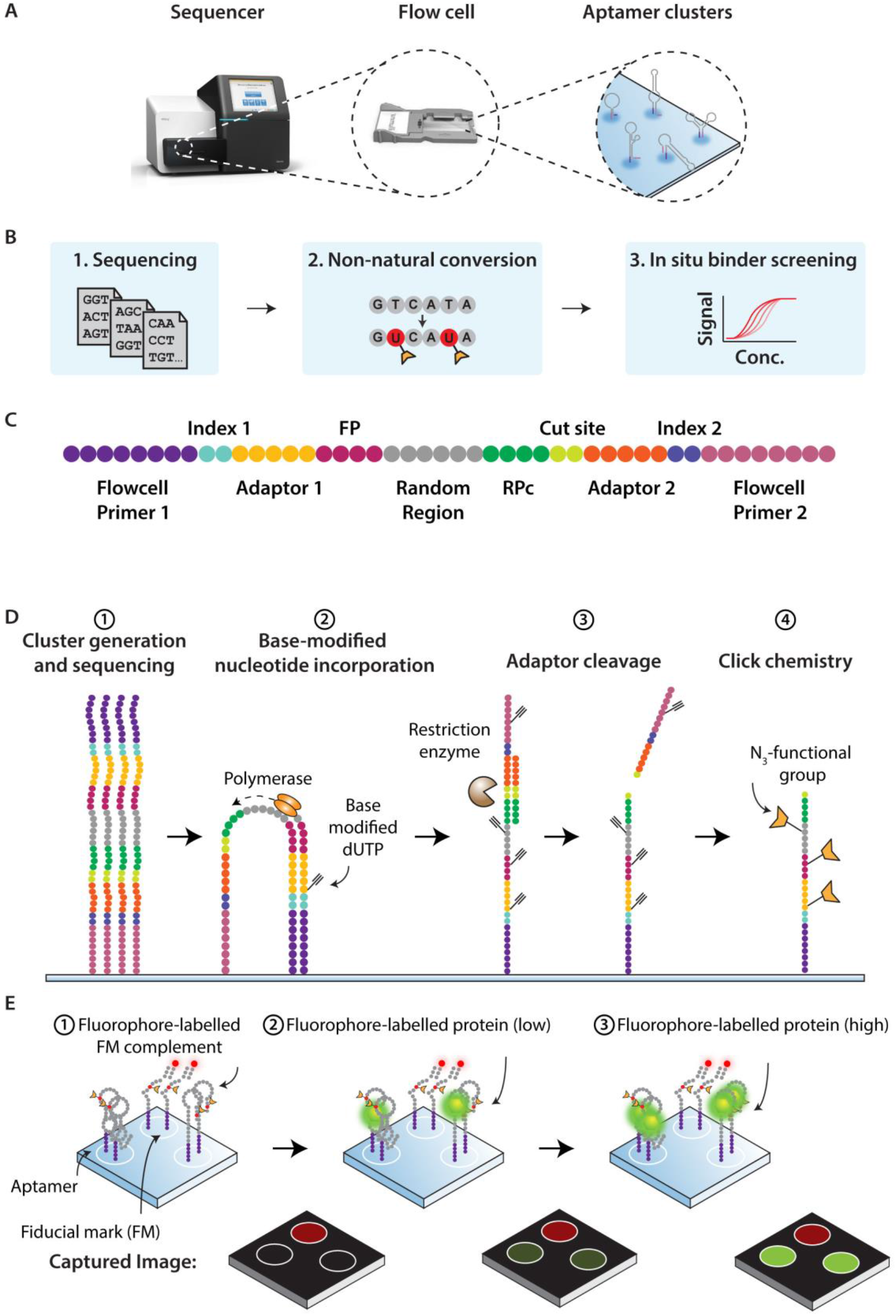
Overview of N2A2. **A**) N2A2 uses a MiSeq flow-cell to generate base-modified aptamer clusters and screen their performance. **B**) The workflow begins with sequencing of a DNA library. During this process, the DNA clusters are converted into base-modified aptamer clusters, which are then incubated with fluorescently-labeled protein to screen for affinity and specificity, with intensity information collected for each cluster. **C**) The initial DNA pool, which comprises a randomized region (gray) flanked by primer-binding sequences (magenta, dark green), is subjected to two amplification steps to introduce Illumina flow-cell primer sites (pink, purple), Illumina ‘adaptor’ sequencing primers (gold, orange), Illumina-defined indices (teal, dark blue), and an EcoRI cut site (light green). **D**) The base-modified aptamer synthesis process. (1) DNA is covalently attached to the flow-cell, natural DNA clusters are formed, and sequencing by synthesis is performed during read 1. (2) During the paired-end turnaround, C8-alkyne dUTP is incorporated into the DNA by KOD-XL polymerase. (3) Flow-cell primer 2, index 2, and adaptor 2 sequences are cleaved from the clusters by EcoRI. (4) An azide-tagged functional group is conjugated to the alkyne handle via click chemistry. **E**) Assessing binding on the flow-cell surface. (1) During read 2, base-modified aptamer clusters are incubated with a fluorescently-labeled sequence complementary to a fiducial mark DNA, which is included in the assay as a positive control, and then imaged. (2) The aptamers are first incubated with a low concentration of fluorescently-labeled protein and imaged, and (3) this process is repeated with higher concentrations of fluorescently-labeled protein.

We first enrich our initial library for binding to the target using conventional enrichment methods (*e*.*g*., SELEX or particle display) to reduce the pool’s diversity to a scale that can effectively be screened on the N2A2 (∼10^6^ sequences).^23,24^ The enriched aptamer pool is then amplified with a pair of adaptor sequences, producing a library of aptamers that feature an EcoRI recognition sequence adjacent to the RPc sequence (**Fig. 1C**). In a separate amplification step, Nextera XT indices that include the Illumina-defined flow-cell primers are added onto the 5’ and 3’ ends of each library molecule. The details of library preparation are provided in the **Supplementary Methods**. All DNA sequences are provided in **Table S1**. Once the library is prepared, we perform paired-end sequencing of the aptamer pool with a V3 MiSeq kit. During this process, the aptamer pool is displayed on the flow-cell as antisense strands, with a 5’ reverse primer sequence and 3’ forward primer complementary sequence. We note that sequencing produces a FASTQ file that represents these sequences as sense strands.

The conversion of DNA into base-modified aptamers is achieved through a four-step process (**Fig. 1D**). The first read of sequencing generates the natural DNA template and FASTQ data (step 1). In the paired-end turnaround step, C8-alkyne-dUTP is substituted for native dTTP during bridge PCR using the KOD-XL polymerase, which can incorporate the modification with high fidelity (step 2).^25,26^ This produces aptamer clusters with alkyne handles that are compatible with post-synthesis modification via click-chemistry reaction. This enables us to replace every T with virtually any chemical functional group that we wish to incorporate. Next, we remove the adaptor and flow-cell primer sequences that were added to the 3’ end of the aptamers during cluster formation in order to prevent potential steric hindrance between the aptamer and the target (step 3). As mentioned above, our library incorporates an EcoRI recognition sequence between the aptamer and the adaptor 2 primer; by hybridizing a complementary strand to the primer and recognition sequence, we form a double-stranded cut-site that enables enzymatic excision of the 3’ sequencing primer, index, and flow-cell primer sequences. Finally, during the second read of the paired-end sequencing process, the desired chemical modifications are conjugated to the aptamer through click chemistry (step 4).

We note that this entire process is performed directly by the MiSeq on the flow-cell, with no human intervention. This is achieved by editing the XML files to control the MiSeq instrument to incorporate the necessary reagents (*i*.*e*., non-natural dNTPs, enzymes, and buffer) stored in the custom tubes in the Illumina sequencing cartridge. Details of the modifications to the XML files are provided in **Supplemental Code**. Only modest hardware modifications are required, as shown in **Fig. S1**. Control experiments for modified cluster generation and click reactions are shown in **Fig. S2 and S3**.

Once the conversion process is complete, we perform affinity screening *in situ* within the flow-cell. A single fiducial mark sequence is included in the sequenced DNA library, and we incubate the flow-cell with a fluorescently-labeled oligonucleotide complementary to this sequence, which serves as a positive binding control in all steps of the experiment (**Fig. 1E**, step 1). Next, we incubate the aptamer clusters on the flow-cell with fluorescently-labeled target proteins (step 2), wash with buffer, and use the imaging system of the MiSeq instrument to capture images of the flow-cell. These images provide the fluorescence intensity of all clusters simultaneously, which is representative of the amount of target captured by each aptamer cluster. Next, we increase the concentration of the target protein, wash, and image, repeating this process at multiple concentrations. In this study, up to seven different concentrations were tested, with additional cycles included to confirm return to baseline. These intensities provide binding information for all modified aptamer clusters on the flow-cell (step 3). As we show below, it is also possible to perform this process in complex matrices such as 1% serum, which allows us to measure the specificity as well as the affinity of individual base-modified aptamers on the flow-cell.

Finally, we have developed custom software tools that create a 1:1 mapping between each aptamer sequence and the fluorescence intensity profile of its cluster across a range of target concentrations. Briefly, we extract the internal binary files produced by the MiSeq software during image analysis. The MiSeq’s internal algorithm performs background normalization and Gaussian intensity fitting to convert raw cluster images to mapped cluster intensities. These are organized into .locs files, which contain the unique physical address of each cluster (expressed as the cluster’s tile on the flow-cell and *x*/*y*-coordinates on that tile), and .cif files, which contain the extracted fluorescence intensity of each cluster in each of the four fluorescence channels. Together, these files provide an intensity map of every cluster on the flow-cell. Next, we extract the tile and *x*/*y*-coordinates for each sequence as provided in the output files using the .fastq format. In this way, we obtain both sequence and location information for each cluster on the flow-cell. Finally, we cross-reference the intensity map from the .locs and .cif files with the sequence map obtained from the .fastq file in order to create a linked 1:1 sequence-intensity map. This yields a direct link between an aptamer’s sequence and its intensity at a given target concentration.

### Identifying optimal base-modifications with N2A2

As an initial demonstration of N2A2, we assessed our platform’s capacity to compare how two different chemical modifications alter the affinity of base-modified aptamers for VEGF. Multiple aptamers have been previously reported for VEGF in the literature^18,27–28^, and the US Food & Drug Administration has approved an RNA aptamer-based drug (Macugen) for age-related macular degeneration that binds to VEGF with picomolar affinity^29^, making VEGF an ideal and well-characterized target to demonstrate the capabilities of N2A2 as an effective platform for the characterization of base-modified aptamers.

We conducted three separate screens, assessing VEGF binding to natural DNA, tyrosine (Y)-, and tryptophan (W)-modified aptamers. We chose Y and W modifications because these amino acids are known to critically contribute to epitope recognition in antibodies^30^ and are five times more likely to be found at binding interfaces between proteins relative to other amino acids.^31^ Additionally, Somalogic has experimentally demonstrated through crystallographic and nuclear magnetic resonance (NMR) spectroscopy data that aptamer modifications incorporating these residues directly interact with protein targets, most likely as a consequence of accessing chemical functionality that is not accessible to the four natural DNA bases alone.^32^ We began by pre-enriching our initial random library (∼1 nanomole) to a scale that could be effectively screened using N2A2. We achieved this by performing a single round of conventional SELEX followed by five rounds of particle display^33^ using natural DNA. The resulting nucleic acid pools were then synthesized and characterized on N2A2 as natural DNA or Y- or W-modified DNA. We introduced fluorescently-labeled VEGF at concentrations ranging from 10 pM to 100 nM into the flow-cell, with a total of 5–7 protein concentrations for each screen. (**SI Methods Section 1.10)**. To minimize the effect of measurement noise, we averaged the fluorescence intensity of aptamers represented by more than 10 clusters with identical sequences on the flow-cell.

Among the published DNA aptamers for VEGF described to date, the SL2B aptamer discovered by the Yung group exhibits the highest affinity.^18^ We used this aptamer as a positive control for our natural DNA N2A2 screen (**Table S1**). We aligned the aptamer sequences from our screen to the intensity data as described above and removed sequences of incorrect lengths or containing other errors (**SI Methods Section 1.13, Table S2**)

Both base-modified aptamer populations showed increased binding to VEGF relative to the natural DNA library, but the W-modified aptamers exhibited considerably higher affinity compared to both natural DNA and Y-modified aptamers. (**Fig. 2A**) We also noted that the SL2B aptamer (dotted red line, top histogram) performs slightly below the average of our natural DNA pool. Many strong signal outliers can be seen for both the W- and Y-modified pools in the boxplots. Based on these findings, we determined that the W modification is a superior choice for the selection of high-affinity VEGF aptamers, and focused our subsequent efforts on the characterization of W-modified aptamers.

**Figure 2:**
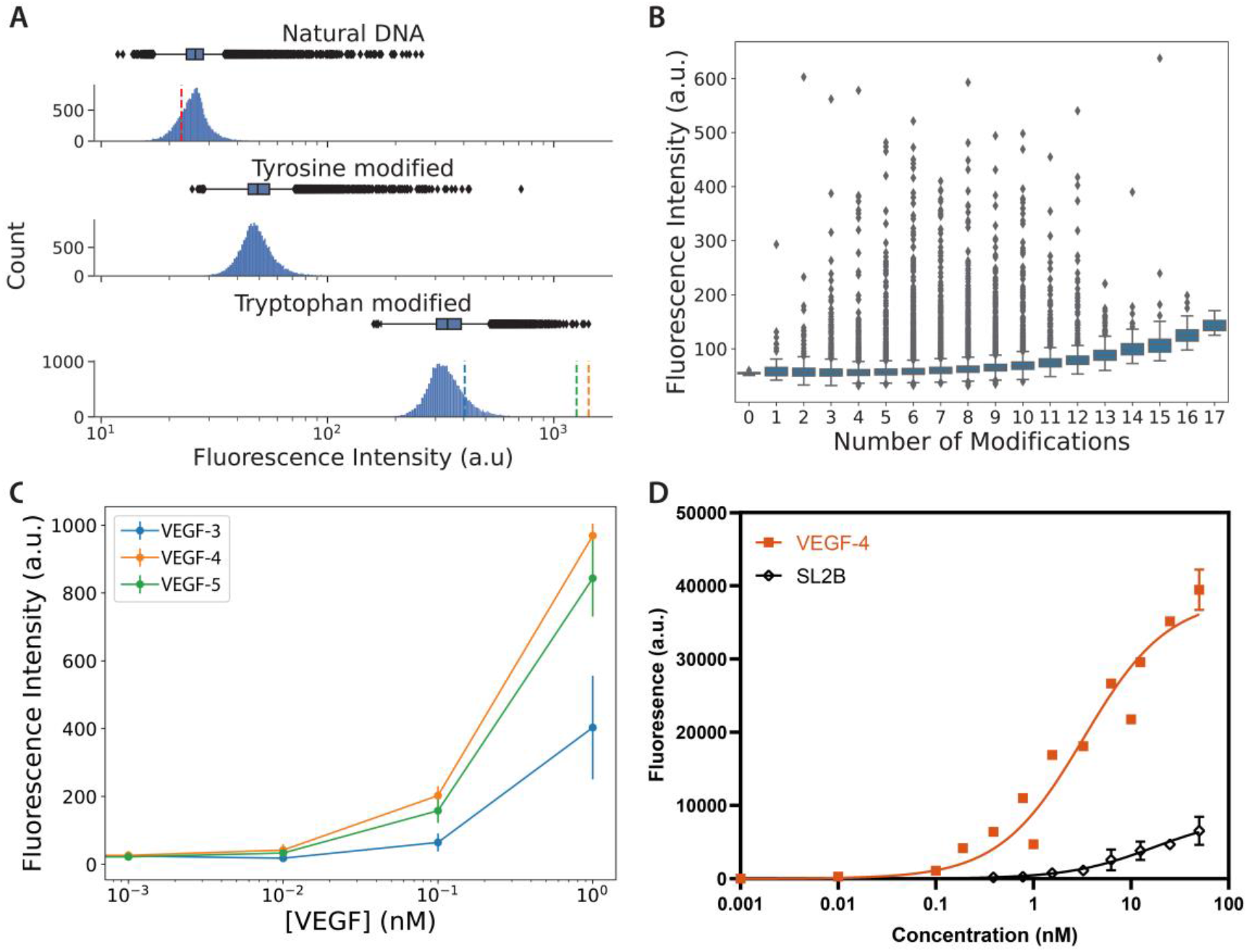
Characterization of base modified VEGF aptamers with N2A2. **A**) Box and whisker plots and histograms of fluorescence intensity for all natural, Y-, and W-modified aptamers at 1 nM VEGF. Histograms show the number of aptamers associated with a given level of fluorescence intensity. The control SL2B aptamer is marked with a dotted red line in the top panel. N2A2-selected aptamers TV3 (blue), TV4 (orange), and TV5 (green) are marked in the bottom histogram. The boxplot is plotted on the same x axis and shows the quartiles of the data set, whiskers representing 1.5x the interquartile range (IQR), and rare high-performing outlier aptamers (diamonds) that are not easily seen on the histogram. **B**) Box and whisker plots of fluorescence intensity at 100 pM VEGF versus the number of W modifications for each W-modified aptamer. Boxes show quartiles of intensity for each aptamer with varying numbers of modifications, and whiskers show 1.5x IQR. Aptamers with intensities greater or less than 1.5 IQR are marked as outliers (diamonds) **C**) Binding curves as measured on the flow-cell for W-modified aptamers VEGF-3, −4, and −5. Error bars represent 1 standard deviation. **D**) Binding curve as measured by flow cytometry for W-modified aptamer VEGF-4 (K_d_ = 3.3nM) and natural DNA aptamer SL2B (K_d_ =16.9 nM) (Background subtracted, fit to specific binding equation, n = 2).

Interestingly, we found that the binding signal of the aptamers was not linearly proportional to the number of W modifications (**Fig. 2B)**. The average fluorescent signal of aptamers increased along with the number of modifications, indicating that the W-modifications positively affect aptamer affinity, but the aptamers with the strongest signal were not the ones with the most modifications. Instead, several of our top binders had only 2–4 W modifications. Additionally, we observed that in the top 1% and 0.1% of binders with the strongest signal, even aptamers with higher numbers of W modifications generally tend not to contain long contiguous stretches of modified bases (**Fig. S4**).

Since N2A2 allows us to characterize many copies of the same sequence, we examined the relationship between copy number and binding performance, as aptamer studies often rely on high copy number in the final round of enrichment to choose potential hits.^24^ We selected a trio of aptamers from the flow-cell: VEGF-3 represented the top of the 3^rd^ quartile of performers, VEGF-4 had low copy number and high binding signal, and VEGF-5 had high copy number and high binding signal (**Fig. 2A, C**). We see that VEGF-4 actually exhibits the best performance of the three aptamers, demonstrating how N2A2 allows us to identify promising low-copy-number hits that would otherwise have been overlooked with purely sequence-count-based SELEX analysis.

We used a bead-based flow cytometry assay^33^ to characterize eight W-modified aptamers (including VEGF-3, −4, and −5) that exhibited target-binding signal in the top quartile of the natural, Y, or W N2A2 experiments at 1 nM VEGF. We found that all eight aptamers selected with N2A2 consistently performed better than SL2B **(Fig. S5, Table S3**). The best aptamer, VEGF-4, showed a K_d_ of 3.3 nM in flow cytometry assays. In comparison, the positive control SL2B aptamer exhibited a K_d_ of 16.9 nM using the same assay. Importantly, the affinity of SL2B as measured on the N2A2 was consistent with measurements obtained with the bead-based flow cytometry assay (**Fig. S6**). We also validated the performance of aptamer VEGF-4 by biolayer interferometry (BLI) and again confirmed that this non-natural aptamer outperforms SL2B in terms of VEGF affinity **(Fig S7**). These results demonstrate how N2A2 experiments can guide the selection of an aptamer with optimal chemical modifications for a given target and facilitate the rapid characterization of many base-modified aptamers in parallel.

### N2A2 enables screening for specificity

N2A2 can also be utilized to simultaneously screen base-modified aptamers for both specificity and affinity. As an example, we isolated a base-modified aptamer that can specifically bind to insulin in diluted human serum. For this experiment, we chose a phenylalanine (F) modification based on the crystal structure of the insulin receptor, which reveals that F plays an important role in the hydrophobic pocket of the insulin-binding site.^30^ As noted above, this residue also tends to be over-represented in protein-binding sites and enhances the binding of base-modified aptamers. We note that insulin is a polypeptide hormone made up of 51 amino acids (molecular weight 5.8 kDa),^34,35^ and generating high-affinity aptamers to polypeptides is much more challenging than for larger proteins. To our knowledge, only one aptamer has been described in the literature for insulin to date (by Yoshida *et al*.), which exhibits a K _d_ of ∼10 μM.^19^

To generate F-modified insulin aptamers, we performed two rounds each of positive and negative SELEX and two rounds of particle display (**SI Methods Section 1.8**), and the resulting pool was characterized on N2A2 (**Table S4**). Briefly, we measured the fluorescence intensity of F-modified aptamer clusters when incubated with 1, 10, and 25 μM insulin in buffer or 1% human serum (**SI Methods Section 1.11; Fig. S8, S9**). By comparing the cluster intensities in buffer and serum, we can readily discriminate between “non-specific” aptamers, which exhibit fluorescence under buffer conditions but no longer generate signal in serum, from “specific” aptamers that retain their fluorescence intensity in both conditions (**Fig. 3A**). For each aptamer cluster, we examined the z-score at each insulin concentration in both buffer and 1% serum to reduce cycle-to-cycle variance and make the populations more directly comparable (**SI Methods Sections 1.15**). Z-score is defined as (x – μ) / σ, where x is the aptamer fluorescence, μ is the mean fluorescence of the entire population, and σ is the standard deviation of the entire population. Since we have many aptamers and aptamer families and want to describe the standard deviations from the population mean, the z-score is an appropriate metric to compare each aptamer to the overall population in a given cycle.

**Figure 3.**
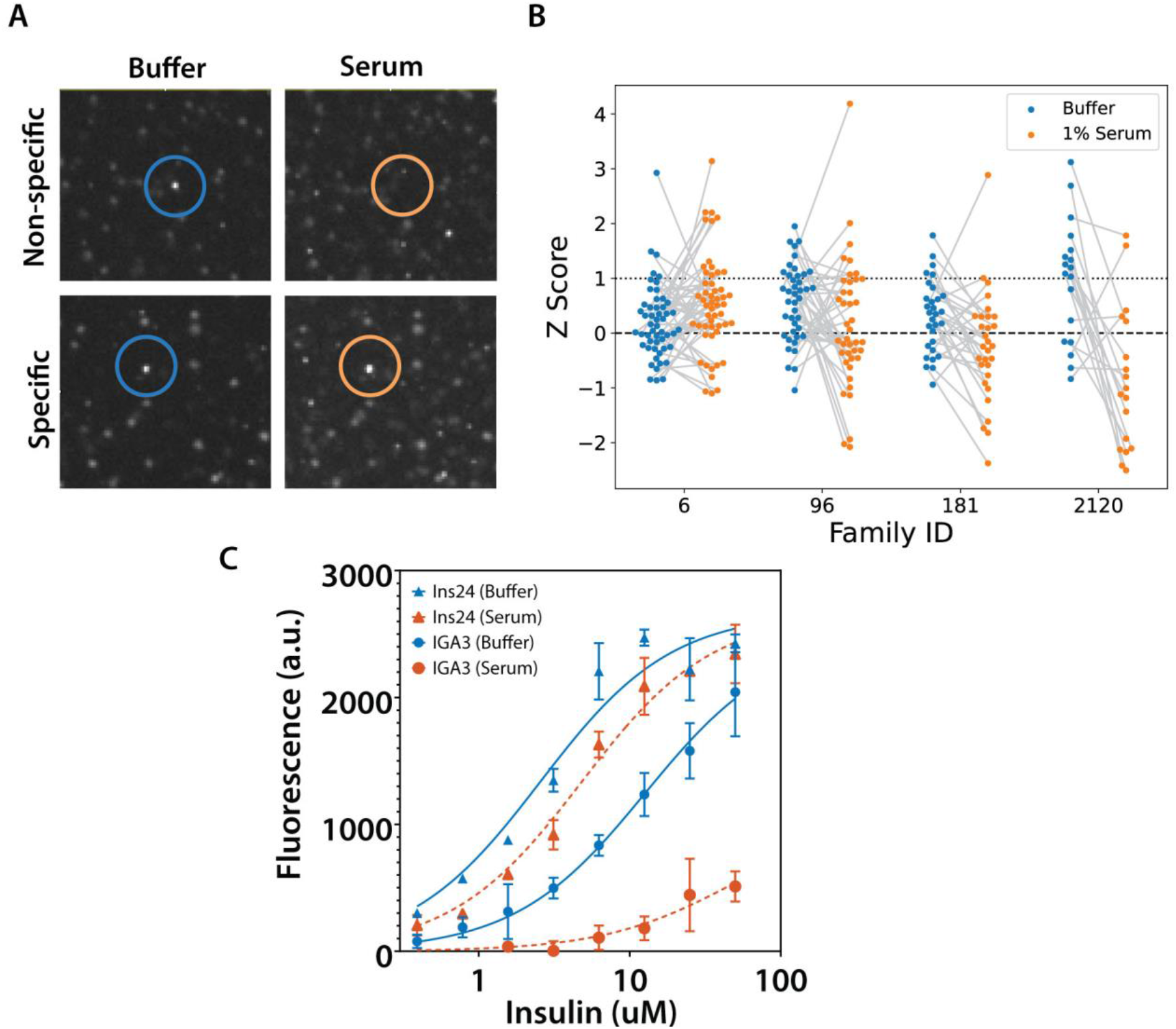
Assessing F-modified insulin aptamer binding in buffer and serum. **A**) N2A2 enables differentiation of target-specific aptamer clusters. A non-specific aptamer (top row) shows a strong signal from insulin binding in buffer (left), but this signal disappears in serum (right). In contrast, a high-specificity aptamer (bottom row) shows strong insulin binding in both buffer and serum. **B**) Z-scores from four representative families: 6 and 96, which are largely specific and retain binding in both buffer and 1% serum, and 181 and 2120, which generally show reduced binding in 1% serum. The z-scores of each individual aptamer are plotted in both buffer and serum conditions, with grey lines connecting datapoints for each sequence. A Z score of 1 is 1 standard deviation above the mean. **C**) A flow cytometry-based bead-binding assay for ins24 and natural DNA aptamer IGA3 in buffer (blue) and 1% human serum (orange).

Finally, N2A2 allows us to examine both individual aptamers and aptamer family performance. Family clustering has previously been used to choose SELEX aptamer candidates after HTS^36^, however, N2A2 allows us to visualize how an entire family performs and which are the best candidates within the family. We performed family clustering with a Levenshtein edit distance of ≤5 (**SI Methods Section 1.14; Fig. S10)** and identified two distinct types of aptamer families. One group of families generally showed a reduced z-score in 1% serum, such as families 181 and 2120, while the other mostly retained their fluorescent signal in 1% serum, such as families 6 and 96 (**Fig. 3B)**.

We tested many aptamer candidates from family clusters which retained binding in serum, as well as aptamers which did not cluster into a family but had high binding performance, at various insulin concentrations by flow cytometry **(Table S5; S11)**. Family 6 stood out because its mean fluorescence was the highest among all families, and we further investigated sequence ins24—a high copy-number representative. We used the same bead-based flow cytometry method mentioned above to compare the insulin affinities of ins24 and IGA3, the Yoshida *et al*. aptamer^19^, and obtained K_d_s of 2.6 ± 0.6 μM and 12.7 ± 1.1 μM, respectively, in buffer (**Fig. 3C**). Critically, only ins24 retained equally strong affinity when we performed the same assay in 1% serum, with a K_d_ of 4.8 μM ± 0.7. In contrast, IGA3 exhibited dramatically lower affinity, and we were unable to obtain a meaningful K_d_ measurement. A follow up microscale thermophoresis experiment with fluorophore-labeled ins24 and unlabeled insulin confirmed that our aptamer has a K_d_ of 2.4 ± 0.4 μM and 2.6 ± 0.6 μM in buffer and serum conditions, respectively (**Fig. S12**). Although the affinity of the selected aptamer is modest, these results demonstrate that N2A2 can rapidly identify base-modified aptamer sequences that outperform natural DNA in terms of binding to challenging small polypeptide targets, and which maintain consistent binding performance even in high-background serum samples.

## Discussion

We have demonstrated here a workflow for the efficient screening and characterization of millions of base-modified aptamers. Our N2A2 platform is designed around a modified Illumina MiSeq instrument and exploits that technology’s sophisticated flow-cell and imaging apparatus to analyze vast arrays of base-modified aptamer candidates *in situ*, in a manner that directly couples binding data to defined sequences based on their position within the flow-cell. Since our approach uses a click chemistry-based modification strategy, one can employ commercially available polymerase enzymes to incorporate virtually any base modification without meaningfully changing the experimental workflow.

As an initial demonstration, we showed that N2A2 can be used to compare the affinity of DNA aptamer libraries incorporating different base modifications and identify which modification confers superior affinity for the target protein VEGF. Our results show that tryptophan modification greatly enhances aptamer affinity for this target, and we subsequently isolated a novel W-modified aptamer with superior affinity to the best natural DNA aptamer in the literature. N2A2 can also be used to simultaneously screen base-modified aptamers for specificity as well as affinity. We screened a library of F-modified aptamers for insulin in both buffer and diluted serum and obtained an aptamer that retained its affinity in both conditions. In contrast, a previously published aptamer with comparable affinity in buffer essentially loses its capacity to bind insulin in serum.

We would like to emphasize that N2A2 is not a replacement for the entire SELEX process; rather, when paired with a few rounds of a well-designed enrichment protocol, N2A2 allows for the simultaneous characterization of a partially-enriched library of ∼10^6^ sequences. This represents a multiple log increase in the number of natural or base-modified aptamers that can be analyzed in a single experiment. Identification of optimal base modifications and sequence variants can be accomplished within 2-3 N2A2 runs. Additionally, due to the versatility of the platform, one can readily test a wide range of experimental conditions—including diverse buffers, complex backgrounds, pH levels—in rapid succession.

This is also a platform with considerable potential for extension in the future—for example, studying single and double mutant libraries of an individual sequence to create sequence fitness landscapes. The scale of this assay is well suited for creating sequence fitness landscapes, as Knight and coworkers did with the Closed Loop Aptameric Directed Evolution (CLADE) approach^37^ and for predicting novel aptamers with machine learning.^38^ Such gain- and loss-of-function mutational landscapes could be deeply informative for assay design or the introduction of further modifications to base-modified aptamers, and could even yield aptamers with further optimized binding properties. Libraries featuring multiple modifications in parallel could be screened using the selection techniques developed by Gawande *et al*^7^. The platform could also be extended to RNA aptamers—previous work from the Lis^15^ and Greenleaf^16^ labs has demonstrated the profiling of RNA-protein interactions on a sequencing flow-cell, where RNA strands were synthesized on the flow-cell after sequencing was performed and the flow-cell was removed from the sequencer. These experiments were conducted with a GAII flow-cell, but the same methodology could readily be adapted to the MiSeq. Alternatively, the range of functions or activities^39^ screened—such as structure switching^40^—could be expanded by modifying the scripting of reagents delivered during read 2. Additional modifications to the XML and driving scripts could also facilitate stepwise sequencing, natural aptamer screening, click modification, and base-modified aptamer screening within a single experiment.

Given that our system is built upon a widely-available sequencing instrument and uses a straightforward and versatile click chemistry approach to achieve aptamer modifications, we believe that N2A2 offers a broadly accessible solution to researchers looking to develop specialized aptamers for challenging applications.

