## Supplemental Information for "Flow-cell based technology for massively parallel characterization of base-modified DNA aptamers"

### Supplementary Information

#### I. Supplementary Methods

1.1 General methods

All DNA oligonucleotides were purchased from Integrated DNA Technologies (sequences described in **Table S1**). Primers were ordered with standard desalting. PCR templates were ordered with PAGE purification. Other than the exceptions noted below, all commercially available reagents and lab supplies were purchased from Sigma-Aldrich or Thermo Fisher Scientific. VEGF was purchased from R&D Systems. Human serum was purchased from Innovative Research. AlexaFluor-647-labeled human insulin was purchased from Nanocs. MiSeq 150 V3 kits were purchased from Illumina. C8-alkyne-dUTP was purchased from IBA Life Sciences. Tyrosine azide, tryptophan azide and phenylalanine azide were purchased from ChemPeP. EcoRI was purchased from New England Biolabs. KOD-XL was purchased from Millipore Sigma. Unless otherwise specified, all chemical reagents were purchased from Pierce Biotechnology. Flow cytometry assays were performed using a BD Accuri C6 flow cytometer. FACS was performed with a BD FACSAria II.

1.2 Hardware modification of MiSeq

Minor modifications were made to a standard Illumina MiSeq to accommodate the additional reagents we wanted to introduce to the flow-cell during binding experiments. Compatible tubing was purchased from Cole Parmer (Part No: 06407-41), couplers were purchased from IDEX Health & Science (Part No: F-130) and a multiport was obtained from Valco Instruments (Part No: EMHMA-CE, 08T-0479L). A fluidic inlet line was added to port 23 in the instrument's internal multiport (Valco Instruments) (**Figure S1**). This port is used to introduce air during cleaning cycles but is not used during sequencing. The fluidic line is linked to the external multiport. During an N2A2 run, our software modifications trigger the external multiport to switch ports, so that the appropriate reagent is drawn into the flow-cell.

1.3 Software modification of the MiSeq

Two software modifications were made to the MiSeq to perform N2A2 experiments and control the external hardware.

First, we modified the run configuration files to perform a custom set of chemistry steps after sequencing, enabling us to automatically perform custom chemical incorporation and binding assays. The custom chemistry definitions include pumping KOD-XL, EcoRI, or buffer from the additional ports in the MiSeq cartridge; heating the flow-cell; incubating reagents on the flow-cell; and pumping in externally stored protein dilutions through the external multiport. Detailed changes to the code can be found in Supplementary Code.

Second, to control our external multiport, we used free automatic file management software (Folder Agent, created by Tony Edgecombe), which monitors the temporary experiment folder and triggers commands upon the creation of .cif files as they are processed. Support for Folder Agent has been discontinued but several alternatives exist (https://alternativeto.net/software/folder-agent/). During the second read of the first cycle, the EcoRI site is cleaved. The flow-cell is then washed with buffer followed by 50 mM NaOH and 0.25% SDS. Depending on whether we are performing a base-modified or natural DNA run, the flow-cell is incubated with click chemistry reagents (see section 1.4) or selection buffer (see section 1.7 for recipe), respectively, for 40 minutes, then washed again. Decision Tree A switches the multiport to the NaOH port after the buffer wash. During the second read of the second cycle, which is the first *in situ* binding measurement, Decision Tree B commands the multiport to switch to the NaOH port during the wash step, start a timer, display "Target Incubation", check the cycle number, and change the multiport to the correct target port. The cycle number and target ports are customized for each individual experiment.

#### 1.4 CuAAC chemistry on the MiSeq

We utilized the paired-end turnaround procedure of the standard MiSeq workflow to introduce the new nucleobases that allow us to convert natural DNA into base-modified aptamers on the flow-cell (**Figure 1B**). We introduced a commercially available polymerase, KOD-XL, and a new dNTP mix, including C8-alkyne-dUTP, into user-replaceable reagent tubes in the MiSeq reagent cartridge. The new PCR mix consists of 804 µl water, 100 µl 10x KOD buffer, 20 µl 10 mM dATP, 20 µl 10 mM dGTP, 20 µl 10 mM dCTP, 20 µl 10mM C8-alkyne-dUTP, and 16 µl 2.5 U/µl KOD–XL. During the paired-end turnaround, instead of using the standard amplification master mix, the sequence recipe was edited to use the KOD-XL PCR mix, which was stored in the reagent cartridge.

During the click chemistry step, reagents were prepared 30 minutes before the MiSeq was commanded to draw the sample. A pre-prepared mixture of 0.1 M $CuSO_4$/0.2 M tris(3-hydroxypropyltriazolylmethyl)amine (THPTA) in $H_2O$ and 10 mM azide-modified tryptophan, tyrosine, or phenylalanine was degassed with $N_2$ for at least 15 minutes. 20 mg of sodium ascorbate was measured out and resuspended in 1 mL $H_2O$ directly before the command. 25 μl of this sodium ascorbate solution was added to the click reaction, and the tube was immediately attached to the external port of the MiSeq. Final concentrations were 1.5 mM $CuSO_4$/THPTA, 0.2 mM azide-conjugate, and 5 mM sodium ascorbate. The alkynes present in the oligos on the flow-cell surface, which had been incorporated by the KOD-XL PCR mix, were the limiting reagent.

1.5 Validating conversion to base-modified aptamers

To validate the complete generation of base-modified DNA clusters, we tested whether our fiducial mark sequence was synthesized in the sense direction. The fiducial mark sequence was indexed with Illumina indices separate from the aptamer pools. We removed the flow-cell from the instrument after the paired-end turnaround and incubated it with 100 μl of 100 nM fluorophore-labeled fiducial mark complementary strand (**Table S1**). If the turnaround was unsuccessful, the fiducial mark would not appear on the flow-cell (**Figure S2A**). Our test showed successful C8-alkyne-dUTP incorporation. New functional groups were introduced to our base-modified DNA strands through the copper click reaction described above, and to validate our reaction procedure on the flow-cell, we introduced an azide-tagged Cy3 label. The appearance of clusters during the second read indicated that our reaction was successful (**Figure S2B**).

1.6 Protein labeling

500 μg VEGF was resuspended in 0.05 M sodium borate buffer and incubated with DyLight 650 N-hydroxysuccinimide (NHS) ester (Thermo Fisher Scientific) for 1 hour at room temperature. This fluorophore was chosen because it aligns with the "A" and "C" imaging channels on the MiSeq. Non-reacted reagent was removed by dialysis (Slide A Lyzer MINI 10K, Thermo Fisher), and the degree of protein labeling was determined using a NanoDrop spectrophotometer. Insulin was purchased pre-labeled with AF647 or biotin.

1.7 SELEX and particle display for VEGF pre-enrichment

One round of regular SELEX was performed with VEGF on particles. 60 μl VEGF (1 μg/μl) was immobilized onto 100 μg Dynabeads M-270 carboxylic acid beads (Thermo Fisher Scientific) according to the manufacturer's protocol for two-step coating using NHS. 6 nmol random library (Primers: **V FP**, **V RP**, **Table S1**) was incubated with 100 µl VEGF-conjugated beads and 100 µl selection buffer (20 mM Tris, pH 7.5, 100 mM NaCl, 5 mM KCl, 2 mM $MgCl_2$, 1 mM $CaCl_2$, 0.1% Tween 20) for 30 minutes at room temperature with rotation. Beads were washed once with 200 µl selection buffer. DNA was eluted from the beads with 200 µl 0.5 M NaOH. Beads were washed once more with 50 µl 0.1 M NaOH. We then adjusted the pH with 25 µl 3M NaOAc, then purified the DNA with a Qiagen MiniElute cleanup kit. The DNA was then amplified for use in particle display. PCR conditions were 95 °C for 2 min; 25 cycles of 95 °C for 15s, 58 °C for 30s, 72 °C for 1 min; and 72 °C for 5 min. The PCR reaction contained 50 µl 2x GoTaq PCR mix, 1 µl 100 µM FP, 1 µl 100 µM RP, 4 µl template DNA, and 44 µl $H_2O$. To determine the correct number of cycles for amplification, we performed a pilot PCR. 5 µl of the reaction was removed at cycles 12, 16, 20, and 24, and then run on a 10% TBE gel at 170 V for 40 min. The cycle that yielded a product of the correct length without forming undesired products was chosen for the final PCR reaction. Five rounds of particle display[1] were performed to further enhance the affinity properties of the library. A binding assay was conducted before every round of particle display to determine the VEGF concentration which would give the highest-affinity binding aptamers, while also adjusting sort gates for stringency and purity. The VEGF concentrations selected for R1-R5 were 30 nM, 60 nM, 20 nM, 25 nM, and 20 nM, respectively.

##### 1.8 SELEX for insulin pre-enrichment

Each round of positive SELEX was performed with bead-immobilized insulin, followed by one round of negative SELEX performed with streptavidin beads. This process was repeated once, for a total of four rounds of SELEX. For positive SELEX, 20 µg biotinylated insulin (Eagle Biosciences) was conjugated to 100 µl MyOne SA C1 Dynabeads according to the manufacturer's protocol in 1X PBS (pH 7.4). 6 nmol random library (Primers: **I FP**, **I RP**, **Table S1**) was incubated with 100 µl insulin-conjugated beads and 100 µl selection buffer for 30 minutes at room temperature with rotation. Beads were washed once with 200 µl selection buffer. DNA was eluted from the beads with 200 µl 0.5 M NaOH. Beads were washed once more with 50 µl 0.1 M NaOH. DNA was recovered from NaOH by adjusting the pH with 25 µl 3M NaOAc, then purified with a Qiagen MiniElute cleanup kit. The eluted DNA was then PCR amplified in the following reaction conditions: 25 µl 2X GoTaq PCR mix, 300 nM **I FP**, 300 nM biotinylated **I RP**, 5 µl DNA, and $H_2O$ up to a final reaction volume of 50 µl. PCR conditions were 95 °C for 2 min; 20 cycles of 95

°C for 15 s, 53 °C for 30 s, 72 °C for 30 s; and 72 °C for 5 min. The amplified DNA was purified with a Qiagen MiniElute cleanup kit and eluted in 20 µl elution buffer (Qiagen). To generate single-stranded DNA, the biotinylated double-stranded DNA was immobilized onto 100 µl SA C1 Dynabeads according to the manufacturer's protocol in 1X Binding and Washing buffer (5 mM Tris-HCl (pH 7.5), 0.5 mM EDTA, 1 M NaCl) in a 1.5 mL Eppendorf tube. The DNA immobilized beads were incubated with 100 µl freshly prepared 0.5 M NaOH for 10 minutes. The tube was placed on a magnetic rack for 2 minutes and the supernatant was collected. Beads were washed once more with 50 µl 0.1 M NaOH. DNA was recovered from NaOH by adjusting the pH with 25 µl 3M NaOAc, then purified with a Qiagen MiniElute cleanup kit and eluted in 20 µl elution buffer.

For negative SELEX, the entire pool of single-stranded DNA was incubated with 100 µl MyOne SA C1 Dynabeads for 30 minutes at room temperature with rotation. We placed the tube in a magnetic separator for 1 min and removed the supernatant. We amplified the DNA in the supernatant according to the same PCR conditions as above.

Two rounds of particle display[1] were performed to further enhance the affinity of the library. We employed 500 nM insulin for both rounds of FACS sorting. Finally, we amplified the resulting DNA pool in preparation for high-throughput sequencing.

1.9 DNA preparation for high-throughput sequencing

Following the Illumina library preparation protocol (https://support.illumina.com/downloads/16s_metagenomic_sequencing_library_preparation.html), we prepared the pools of DNA from our pre-enriched VEGF or insulin libraries for sequencing and binding analysis with N2A2. First, we attached sequencing adaptors (**Table S1).** The adaptor PCR reaction consisted of 5 µl 5 ng/µl pre-enriched library DNA, 5 µl 1 µM FP Adaptor 1, 5 µl 1 µM RP Adaptor 2, 25 µl 2X GoTaq HotStart MasterMix, and 10 µl PCR-grade water. PCR conditions were 95 °C for 2 min; 4-8 cycles of 95 °C for 30s, 55 °C for 30s, 72 °C for 30; and 72 °C for 5 min, after which the desired products were separated on a 10% TBE gel at 170V for 40 minutes, and the correct length product was cut out of the gel and eluted in TE. A pilot PCR was run to determine the correct number of cycles for amplification. 5 µl of the reaction was removed at cycles 2, 4, 6, and 8, and then run on a 10% TBE gel at 170 V for 40 min. The cycle that yielded a product of the correct length without forming undesired products was chosen for the final PCR reaction. Next, we attached Nextera XT indices in a PCR reaction consisting of 5 µl 5 ng/µl DNA,

5 µl Nextera XT Index Primer 1 (N7xx), 5 µl Nextera XT Index Primer 2 (S5xx), 25 µl 2X GoTaq HotStart MasterMix and 10 µl PCR-grade water. PCR conditions were the same as used for the adaptor addition, and products were again gel-purified. DNA was quantified via Qubit or qPCR.

1.10 N2A2 VEGF experimental procedure

The same input DNA stocks and labeled VEGF protein were used for all three experiments, but new sequencing cartridges and flow-cells were used for each experiment. For the natural DNA experiment, natural dNTPs were used during the paired-end turnaround, but the sequence was still cleaved at the EcoRI cleavage site before introducing DyLight-labeled VEGF. During the conversion process, tyrosine or tryptophan azide was clicked onto the alkyne handle and then the sequence was cleaved. After collecting the sequence information, we exposed the aptamer libraries to DyLight650-labeled VEGF at concentrations of 1 pM–100 nM in selection buffer.

Our first tryptophan-modified DNA experiment, which used similar concentrations of VEGF to those used for natural and tyrosine-modified DNA (1–100 nM), was oversaturated with signal. We repeated the experiment and reduced the VEGF concentration range to 1 pM–1 nM. Final protein titrations for each experiment were: natural DNA: 0, 10 pM, 100 pM, 1 nM, 10 nM, 50 nM, 100 nM; tyrosine-modified: 0, 10 pM, 100 pM, 1 nM, 10 nM, 50 nM; tryptophan-modified: 0, 1 pM, 10 pM, 100 nM, 1 nM

1.11 N2A2 insulin experimental procedure

During the conversion process, phenylalanine azide was clicked onto the alkyne handle and then the sequence was cleaved. During the second read, we exposed the aptamer libraries to AF647-labeled insulin at concentrations of 1, 10, and 25 µM in both selection buffer and 1% human serum. Before, in between, and after the insulin titrations in buffer and serum, we also performed fiducial mark cycles to ensure that the flow-cell was returning to baseline.

1.12 Sequencing data alignment to images

During data processing, sequence locations, cluster intensities and cluster locations were extracted and linked. This was achieved with code written in Python, which will be made available upon request. Briefly, FASTQ files are parsed and separated into tiles, keeping only sequence and x-y position data. Locs files are parsed into x and y positions. Locs are rounded via the input functions, and then converted to FASTQ range via shifting, scaling, and rounding. Cif files are parsed into intensities by channel: A, C, T, G, or custom. Locs and FASTQ positions are cross-correlated, with .cif data and FASTQ sequence linked through locs and FASTQ x-y position.

1.13 Sequence pre-processing and sequence-intensity alignment

Raw sequencing reads were trimmed with a custom Python script, which removed SELEX primers. Next, sequencing data were aligned to intensity information. Aligned sequence-intensity data were filtered to ensure that the random region was of the expected length +/- 2 nt (VEGF library: N30, insulin library: N40). Sequences with intensity values greater than 50,000 a.u. at any measured concentration were dropped, as this was indicative of an integer error by the MiSeq software during background subtraction. To speed up processing and reduce noise, only sequences with >10 reads were retained. Additional filtering was applied such that the binding intensity at 0 nM VEGF had to be < 50 a.u. to eliminate false positives.

1.14 Aptamer family clustering

Aptamers were clustered into families based on Levenshtein edit distance ≤ 5 using the Python package Levenshtein. We chose this distance because the library random region is 40 bases, and we aimed to create families of sequences that were only 12.5% different from the seed sequence. Other aptamer clustering software (such as FASTAptamer[2]) uses similar initial metrics for clustering—*e.g.*, Levenshtein distance 7 for a 70-mer random region, equivalent to 10% difference. Family consensus sequences were chosen in order of read number, such that the highest copy-number sequences were used first.

1.15 Z score calculation

Z score was calculated as $z = \frac{x-\mu}{\sigma}$, where $\mu$ is the population mean, and $\sigma$ is the standard deviation of the population. For each family containing 10 or more sequences, we also calculated the mean z-score.

1.16 Flow cytometry binding assays

We performed PCR with natural DNA aptamer candidates in a 100 µl reaction consisting of 67 µl PCR-grade water, 10 µl 10X KOD-XL DNA polymerase buffer, 2 µl each of dATP, dGTP, dCTP, and C8-alkyne dUTP (10 mM), 0.1 µl 10 µM forward primer, 10 µl 10 µM reverse primer, 2 µl KOD-XL polymerase (2.5 µ/mL), 1 µl 100 pM template DNA, and 1.2 µl forward primer-conjugated magnetic beads (~$10^7$/mL). PCR conditions for the VEGF library: 95 °C for 2 min; 25

cycles of 95 °C for 15s, 58 °C for 30s, 72 °C for 1 min; and 72 °C for 5 min. PCR conditions for the insulin library: 95 °C for 2 min; 25 cycles of 95 °C for 15 s, 53 °C for 30 s, 72 °C for 30 s; and 72 °C for 5 min.

We used click chemistry to introduce tyrosine, tryptophan or phenylalanine modifications. Base-modified aptamer beads were resuspended in 10 µl PBS. 7.5 µl of 0.1 M $CuSO_4$/0.2 M THPTA was diluted into 47.5 µl water. 10 µl aptamer beads, 55 µl diluted $CuSO_4$/THPTA, and 10 µl 10 mM tyrosine, tryptophan, or phenylalanine azide were mixed in an amber-colored 2 mL tube. The click reaction was initiated by adding 25 µl 0.1 M sodium ascorbate and vortexing. Sodium ascorbate was prepared immediately before the reaction. The click reaction solution was incubated for 40 minutes under $N_2$, protected from light. Beads were washed twice with 100 µl TE buffer and then resuspended in 100 µL TE buffer.

We then generated monoclonal, single-stranded, base-modified aptamer beads by resuspending the particles in 200 µL 0.1 M NaOH and rotating for 10 min at room temperature. Particles were washed five times with TE buffer and resuspended in 200 µL 10 mM Tris, pH 7.5. Positive control beads with aptamer SL2B (for VEGF) or aptamer IGA3 (for insulin) were generated by ordering the biotinylated aptamer (**Table S1**) and immobilizing onto MyOne SA Dynabeads according to manufacturer's instructions.

1 µl beads and VEGF (0.39–50 nM) or insulin (0.39–50 µM) in 100 µl selection buffer were incubated at room temperature for 40 minutes. The tube was placed on a magnetic rack for 30 sec, washed twice with 100 µl cold selection buffer, and resuspended in 100 µl cold selection buffer. The beads' fluorescence was measured by flow cytometry.

To determine if N2A2 measurements and flow cytometry measurements were comparable for the VEGF experiments, we tested the published VEGF aptamer displayed on beads as well as on N2A2. For the flow cytometry experiments, we measured the fluorescence intensity of aptamer particles incubated in 0.39–100 nM DyLight-labeled VEGF. We normalized the data by subtracting the intensity of the negative control and dividing by the maximum intensity measured. (**Figure S3**)

#### 1.17 Biolayer interferometry (BLI) binding assays

Sequence VEGF-4 modified with 5' biotin-TEG and internal 5-octadiynyl-dU modifications was purchased and HPLC purified by IDT (sequence shown in **Table S3**). 5 nmol of modified VEGF-4 was diluted to a final concentration of 10 µM in 50 µl 10X PBS, 30 µl 0.1 M $CuSO_4$/0.2 M THPTA, 50 µl 10 mM tryptophan azide, and $H_2O$ to a final volume of 450 µl. A 50 mM solution of sodium ascorbate was immediately prepared in $H_2O$, and 50 µl of this was added to the reaction mixture. The solution was degassed using $N_2$ for 5 minutes, then incubated at room temperature for 40 min with rotation. After the reaction was complete, the DNA was purified and exchanged into $H_2O$ using 10K Amicon Ultra-0.5 mL Centrifugal Filter (EMD Millipore) spin columns. This DNA was then diluted to 500 nM in 200 µl selection buffer for BLI experiments. VEGF was diluted to 50 nM–1000 nM in 200 µl selection buffer. Binding assays were performed on an Octet Red 384 instrument with streptavidin (SA) biosensors (Octet Bioforte). 200 µl of 500 nM VEGF-4 was loaded onto SA sensors with response of 0.5 nm, followed by wash steps. Loading time was adjusted to ensure consistent immobilization response between natural aptamers and base-modified aptamers. After 120 s buffer wash, aptamer sensors were dipped into the VEGF solution for 300 s association and then into buffer for 600 s dissociation. Two reference sensors were also prepared; one was dipped into buffer with 0.5 nm aptamer immobilization, while the other was dipped into 1 mM VEGF without any aptamer immobilization. We did not observe any non-specific binding from the reference sensors.

#### 1.18 Microscale thermophoresis (MST) measurements

MST measurements were performed by 2Bind, Germany. 5'-end Cy5-labeled ins24 aptamer with internal C8-alkyne dUTP modifications was prepared by PCR following the same reaction conditions as described in Section 1.17, with 10 µl 10 µM Cy5-labeled forward primer and 10 µl 10 µM biotin-labeled reverse primer. Phenylalanine modifications were attached using click chemistry following the same reaction conditions as Section 1.17. Single-stranded aptamers were generated using streptavidin beads following the manufacturer's protocol. The samples were analyzed on a Monolith NT.115 Pico at 25 °C, with 5% LED power and 40% laser power. A serial dilution of unlabeled insulin was prepared, with concentrations ranging from 122 pM to 4 µM. 5 µl of each dilution step were mixed with 5 µl of 5 nM fluorescently-labeled aptamer. The final reaction mixture was filled in capillaries. No sticking of the target to the capillary walls was observed in the capillary scan, and no sample aggregation or precipitation effects were observed in the normalized fluorescence ($F_{norm}$). $F_{norm}$ values for each MST trace were calculated from the ratio of $F_0/F_1$. Each $F_{norm}$ value was then plotted in a semi-logarithmic manner against the ligand concentration to yield a dose-response curve. See **Fig. S12** for fitting and equation.

#### II. Supplementary Figures

Figure S1: Hardware modification for MiSeq

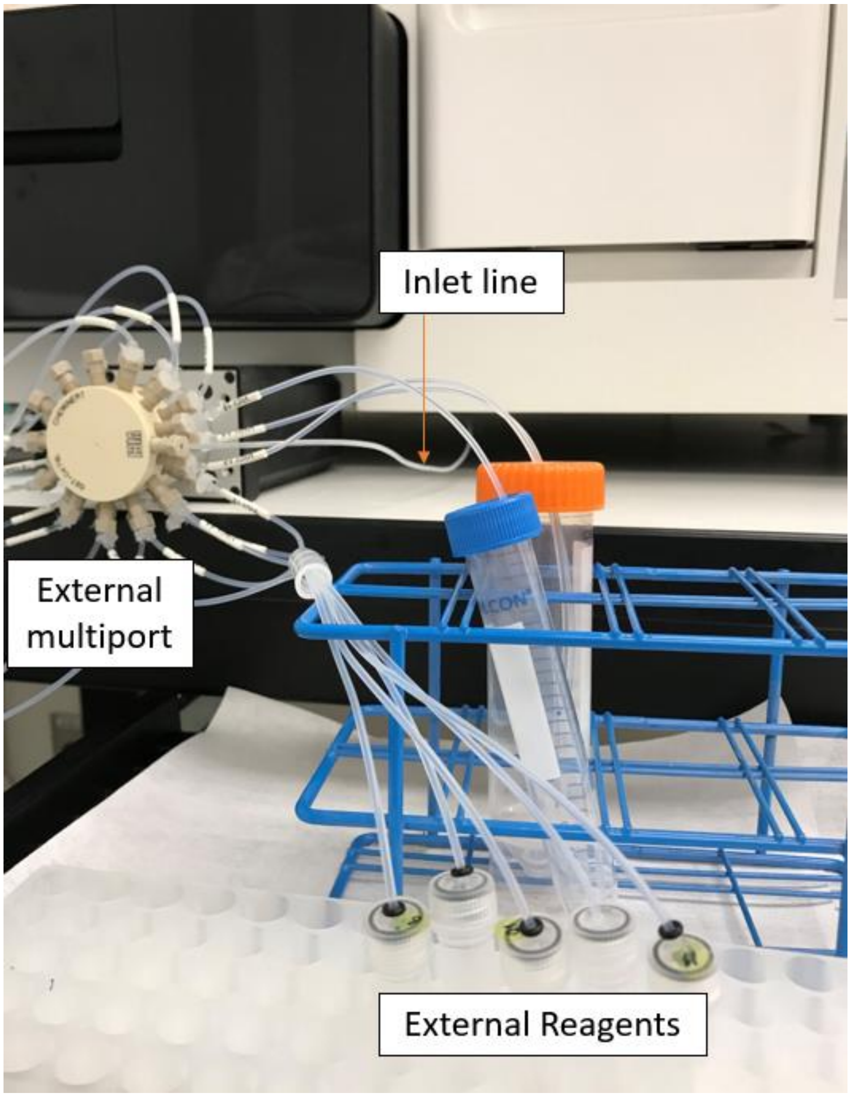


The MiSeq was adapted for N2A2 with a few basic hardware modifications. A single inlet line was connected to line 23 on the internal multiport. This line leads to the external multiport, which can sample from 20 different external reagents.

Figure S2: Validation of the base-modified aptamer generation procedure

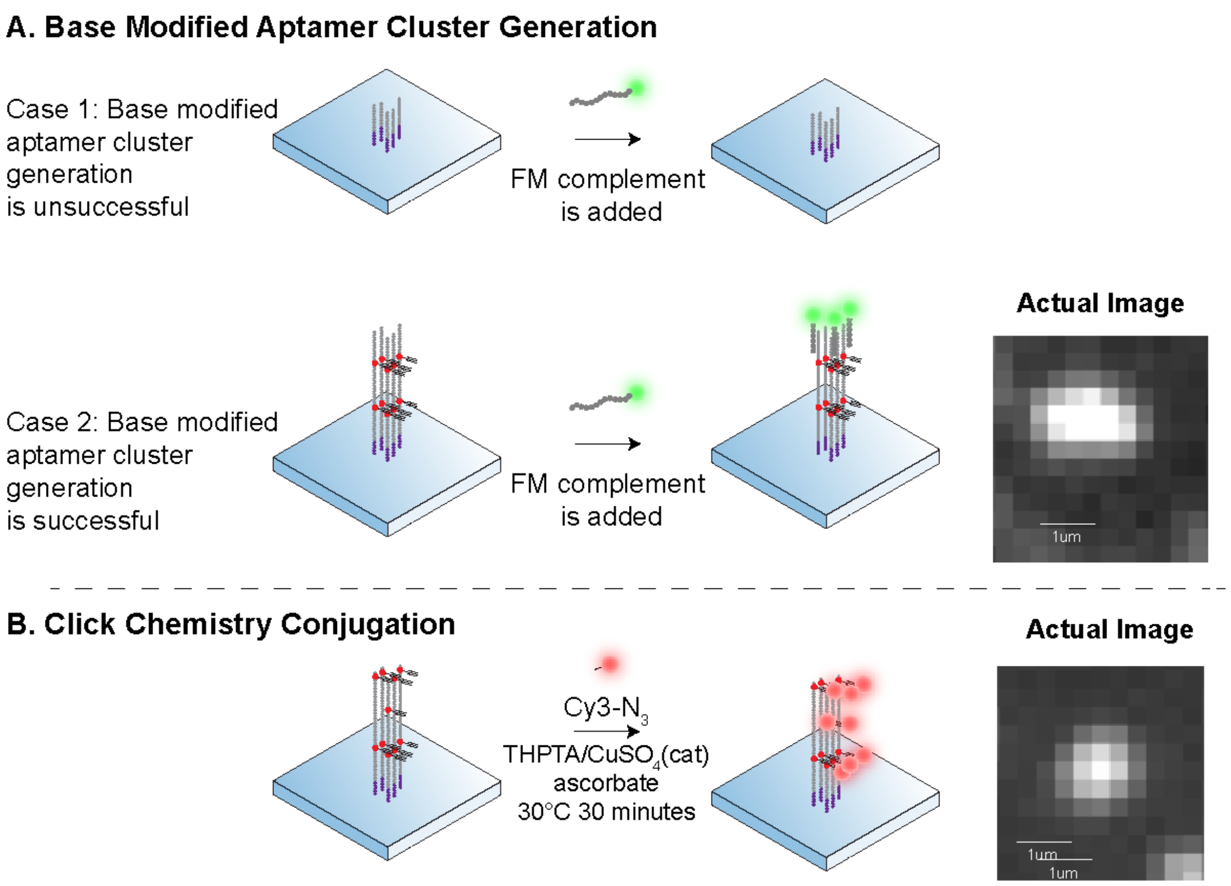


Establishing conditions for base-modified aptamer generation in N2A2. (A) We used fiducial mark (FM) sequences to normalize the background and help with alignment of the clusters. FM sequence hybridization to a fluorescently-tagged FM complement strand demonstrates that all clusters on the flow-cell have successfully incorporated C8-alkyne dUTP. Case 1 and 2 respectively show what occurs if the bridge amplification with C8-alkyne dUTP is unsuccessful or successful, with successful Case 2 image data shown at right. (B) We demonstrated that the optimized click reaction conditions were working by conjugating an azide-tagged Cy3 onto modified dUTPs on the flow-cell.

Figure S3: Click reaction completeness

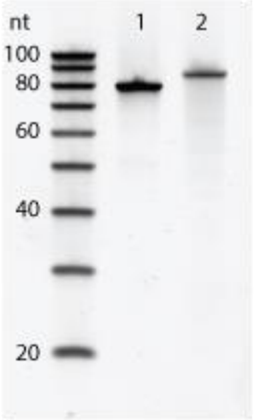


10% TBE urea gel showing alkyne-modified aptamer VEGF-4 generated by solid phase synthesis (lane 1) and alkyne-modified aptamer VEGF-4 with azide-tryptophan modification attached by click chemistry (lane 2). We do not see a secondary band in lane 2, indicating completeness of the click reaction.

Figure S4: Modification profile of W-modified VEGF aptamers

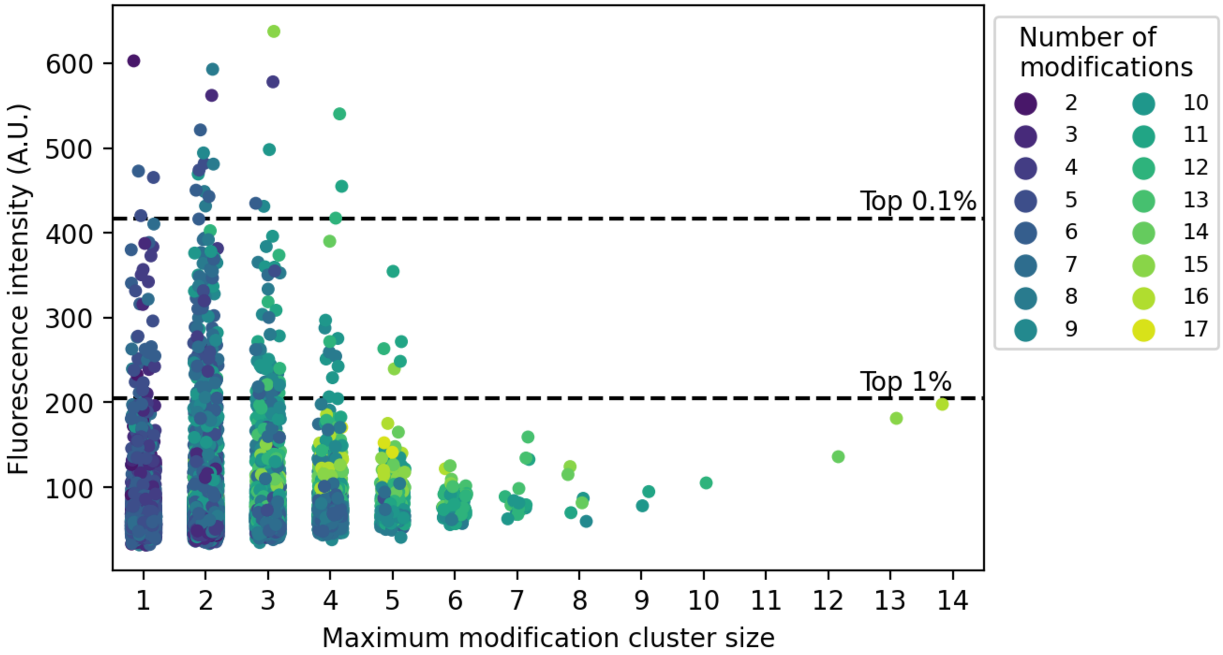


Strip plot showing fluorescence intensity at 100 pM VEGF versus the maximum modification cluster size (defined as the maximum number of adjacent W modifications present in the sequence) for each W-modified aptamer. For example, the sequence 'ACTTGATTTA' would have a maximum modification cluster size of 3, since it contains adjacent T clusters of size 2 and 3. Color indicates the total number of modified bases present in the sequence.

Figure S5: W-modified VEGF aptamers tested by flow cytometry.

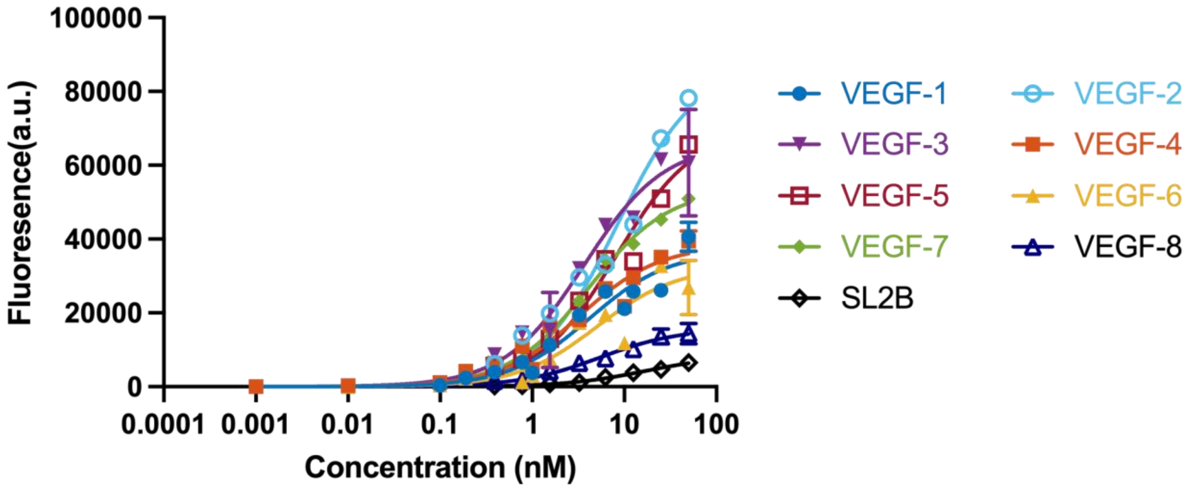


Eight aptamers were chosen for their strong binding performance in natural, tyrosine-, or tryptophan-modified N2A2 experiments for flow cytometry-based binding analysis. Each aptamer was synthesized on beads with the tryptophan modification and measured in a fluorescence-based assay by flow cytometry (n = 1 except for the highest concentration, where n = 2). Background was measured against a titration of blank beads (n = 4). Positive control assays were performed with aptamer SL2B (n = 4). Background was subtracted from all binding curves. One site-specific binding equation was fit to each curve using nonlinear regression. Sequences and $K_d$ are shown in **Table S3**.

Figure S6: Comparison between N2A2 and flow cytometry assessment of VEGF aptamer SL2B

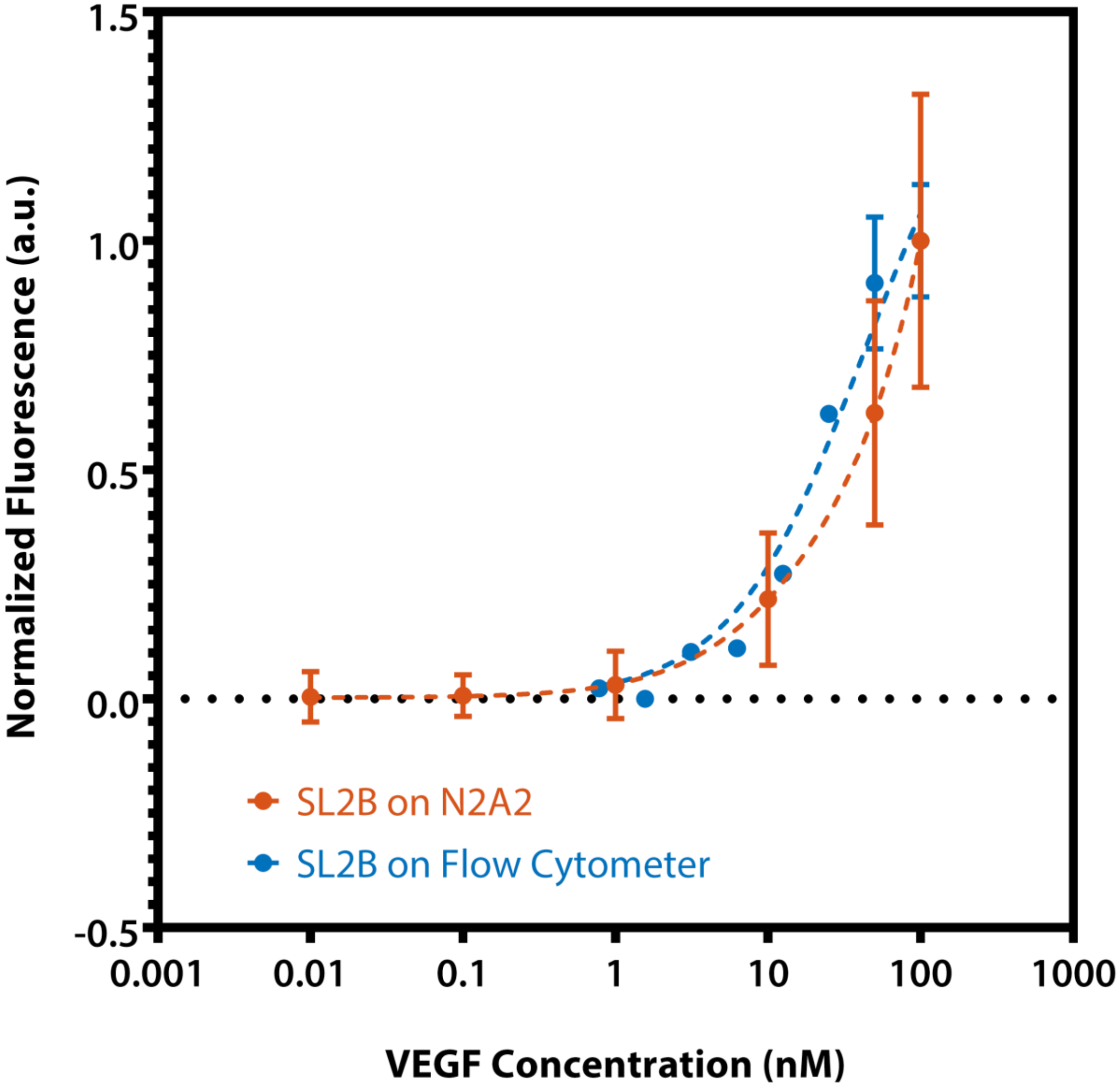


N2A2 data (orange) represents 117 clusters of VEGF aptamer SL2B tested on the flow-cell during the natural DNA VEGF N2A2 run (**Fig. 2A**, top panel, red dotted line). Fluorescence intensities from all clusters were averaged and normalized to a 0 to 1 scale. Data were fitted to total binding (sum of specific binding + background). We also generated SL2B-coated particles and measured VEGF binding by flow cytometry (blue). 10,000 events were collected for each sample, with the background from particle-only measurements subtracted. Data were fitted to the specific binding equation. The two measurement techniques produced similar binding curves, although noise increased as concentration increases.

Figure S7: BLI measurement of VEGF-4 and SL2B

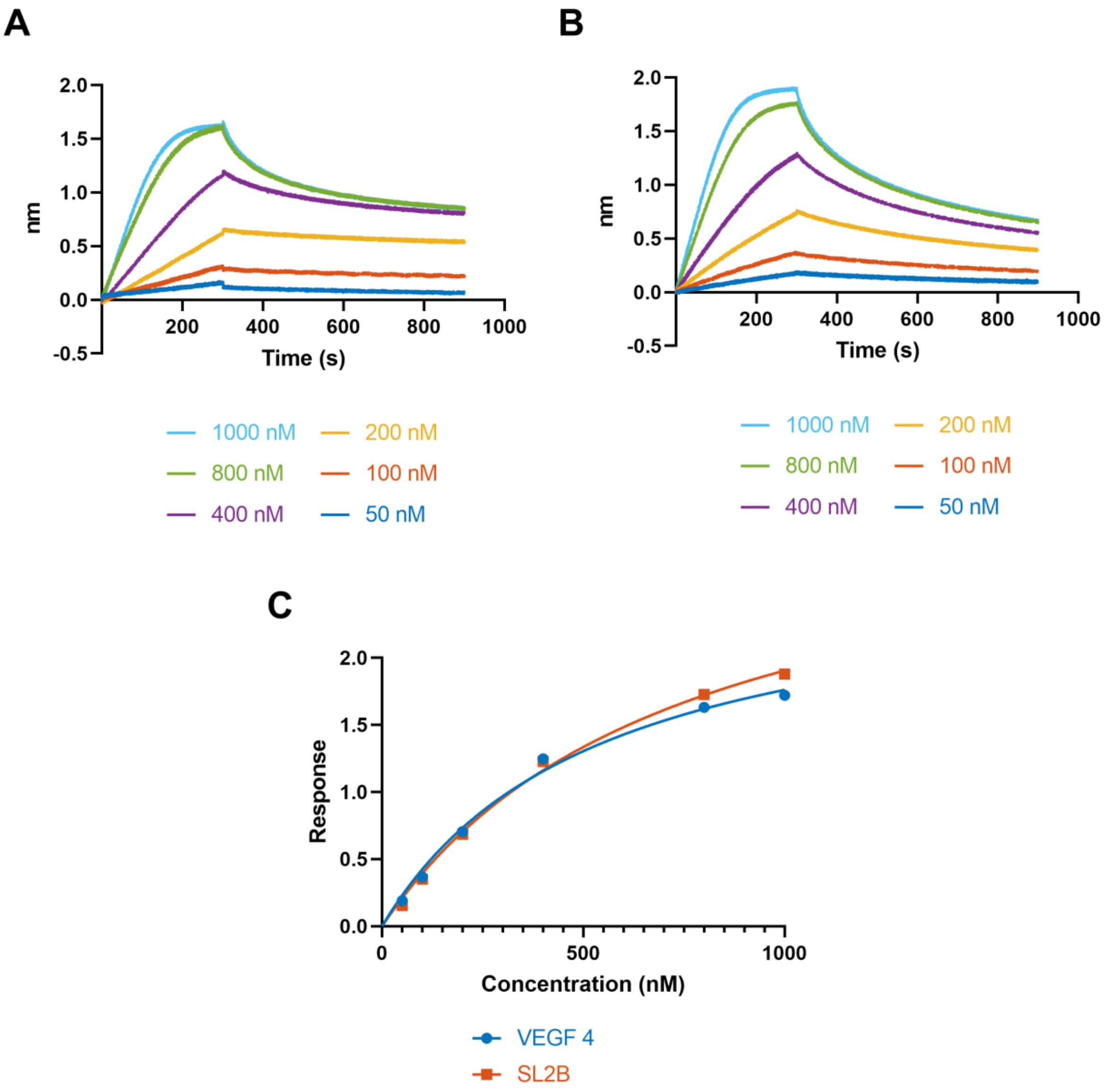


Biolayer interferometry (BLI) measurement of VEGF association and dissociation for A) W-modified aptamer VEGF-4 and B) published aptamer SL2B. C) Steady-state analysis by Octet software calculates $K_D$ based on endpoint values from each association/dissociation curve for titrations of VEGF against SL2B ($K_D$ = 740 nM) and VEGF-4 ($K_D$ = 540 nM), using the following equation: Response = Rmax *Conc/($K_D$+Conc).

Figure S8: Verification of protein stripping

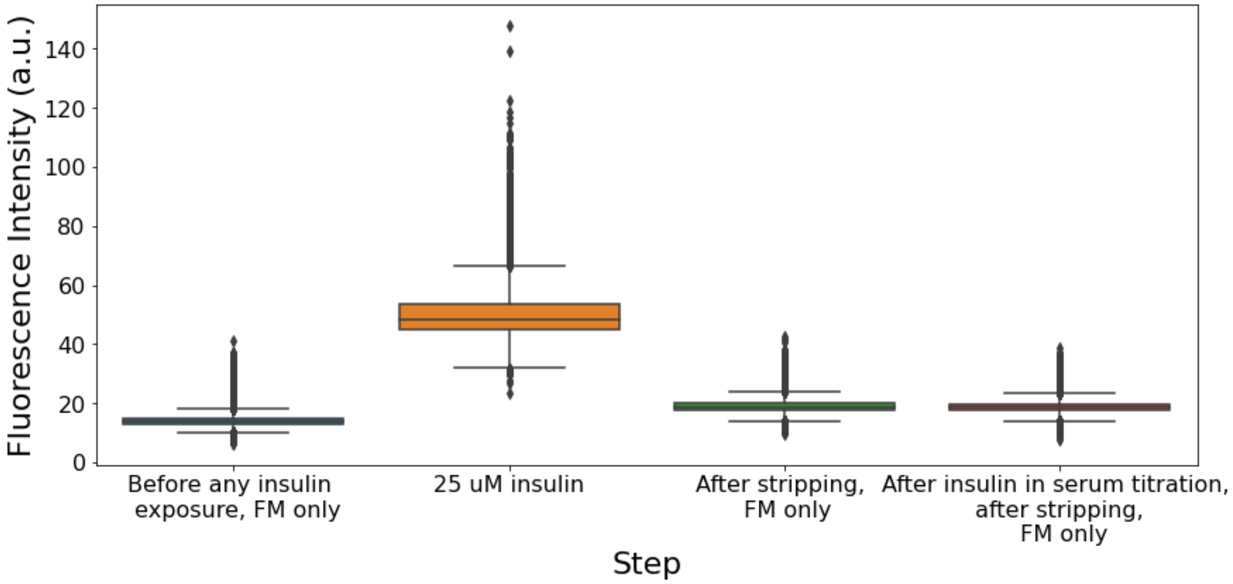


During each protein titration step, the flow-cell is stripped with NaOH to remove any fluorescent signal. This is verified by measuring the fluorescence intensity before exposure to insulin. The labeled fiducial mark complement sequence is provided in the buffer to allow the instrument to focus on clusters, but these are filtered out of this analysis. Box and whisker plots show modified aptamer fluorescence intensity before and after being exposed to labeled insulin, which is then stripped from the surface. After stripping the flow-cell, the fluorescence intensity returns to baseline, almost equivalent to the levels seen before any exposure to insulin.

Figure S9: F-modified insulin aptamers in 1% human serum

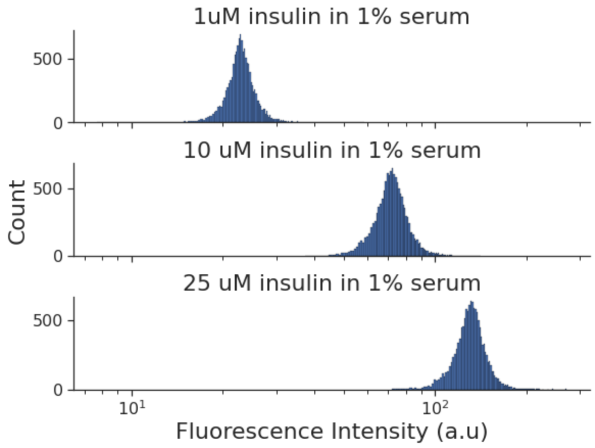


In the insulin specificity experiment, 13,125 unique aptamer sequences (with replicates) were tested for specific binding to 1, 10, and 25 μM labeled insulin in 1% human serum. We observed increased fluorescence intensity as the concentration of insulin increased.

Figure S10: F-modified insulin aptamer families in 1% human serum at 10 μM insulin

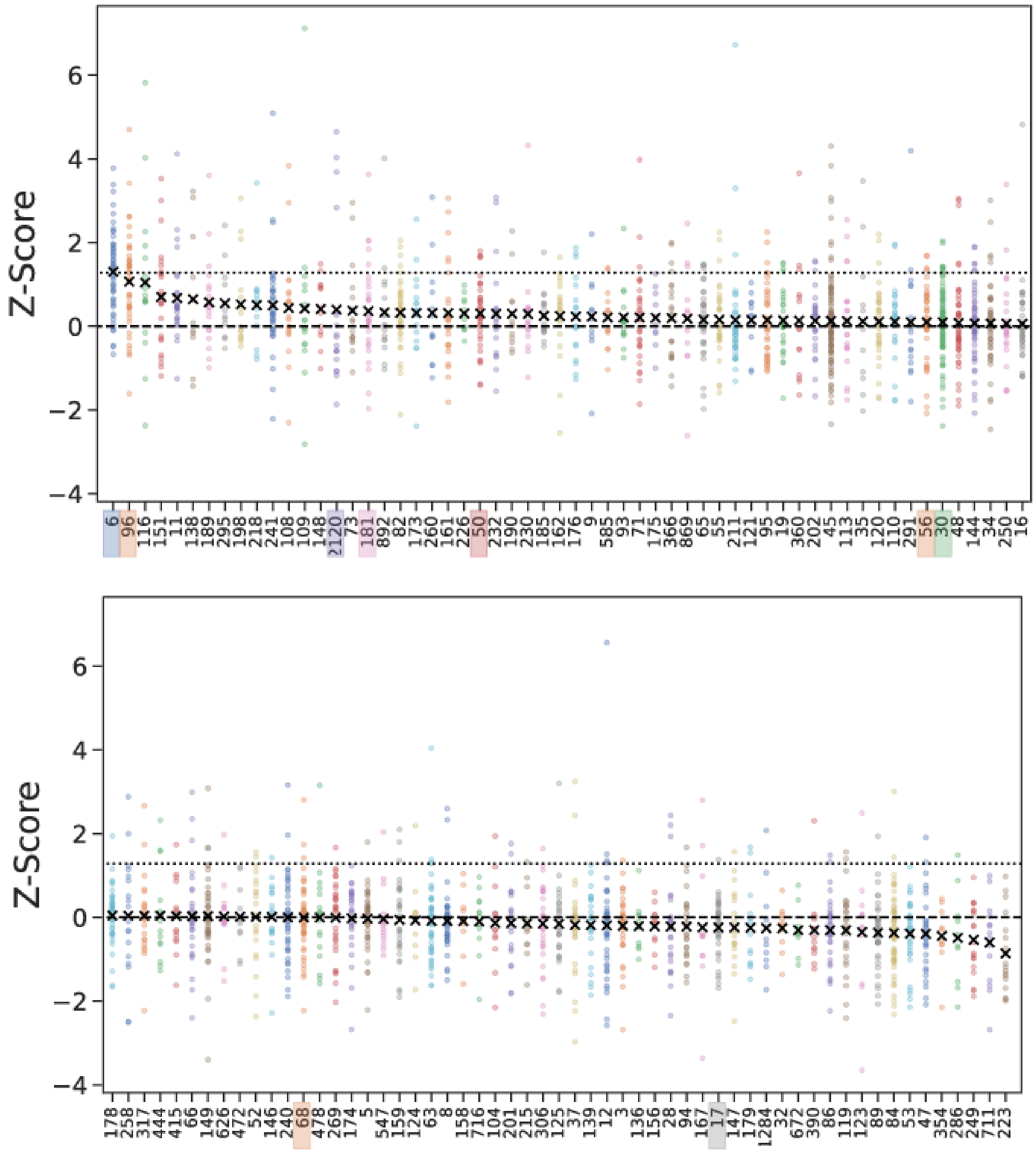


Z-scores were calculated for each aptamer candidate, and aptamers were clustered into families using Levenshtein edit distance ≤ 5. Each color represents an individual family. ‘x’ represents the mean family z-score. The dashed line represents the population mean measured on N2A2. The dotted line represents 90% above the mean, a threshold chosen to down-select the final aptamer families and candidates for testing. Families with 10 or more members are shown, sorted from largest to smallest mean z-scores at 10 μM insulin in 1% human serum. Families tested by bead based flow cytometry are highlighted on the x axis; exact sequences are provided in Table S5.

Figure S11 F- modified insulin aptamer candidates tested by flow cytometry

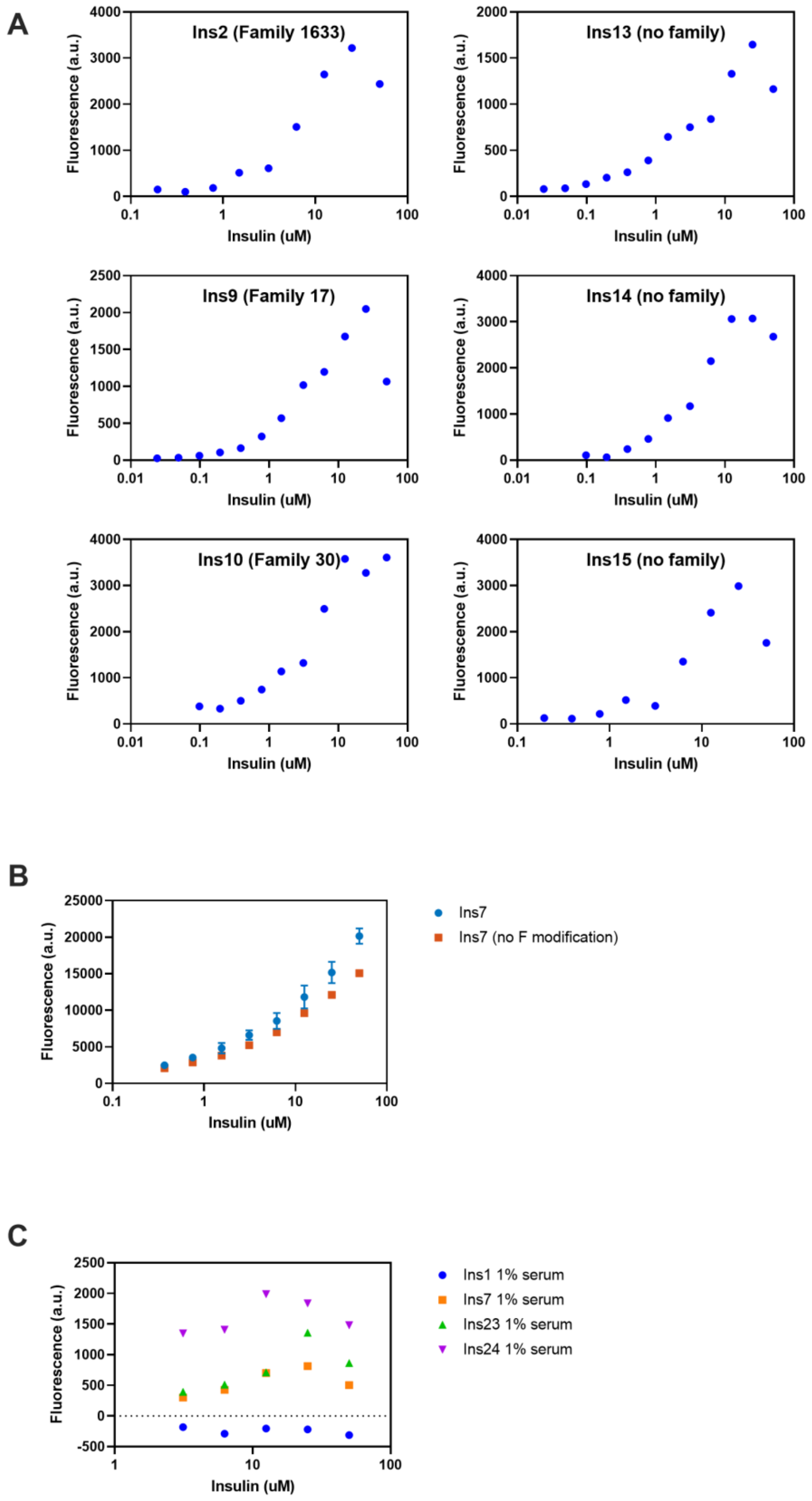

A) Non-natural insulin aptamers identified as binders through N2A2 tested by flow cytometry at various concentrations of insulin in buffer conditions. Some aptamers were chosen by family performance, as shown in Figure S10: Ins2 (family 1633), ins9 (family 17), ins10 (family 30), and other aptamers were chosen by individual performance in N2A2 experiment, although they didn't cluster in a family: ins13 (no family), ins14 (no family), ins15 (no family) B) The addition of the non-natural modification phenylalanine increases the binding performance of an aptamer (blue) compared to the non-modified version (orange). (background not subtracted). Ins7 belongs to family 68. C) Some non-natural insulin aptamers maintain their binding performance in the presence of 1% human serum. Rare sequences with high performance but no family and common sequences, with large families with high performance were both tested: Ins1 (no family), Ins 7 (family 68), ins 23 (no family), ins 24 (family 6) Background was measured against a titration of blank beads and subtracted from all measurements in (A) and (C). All aptamer sequences are listed in Table S5

Figure S12: Microscale thermophoresis validation of Ins24 binding

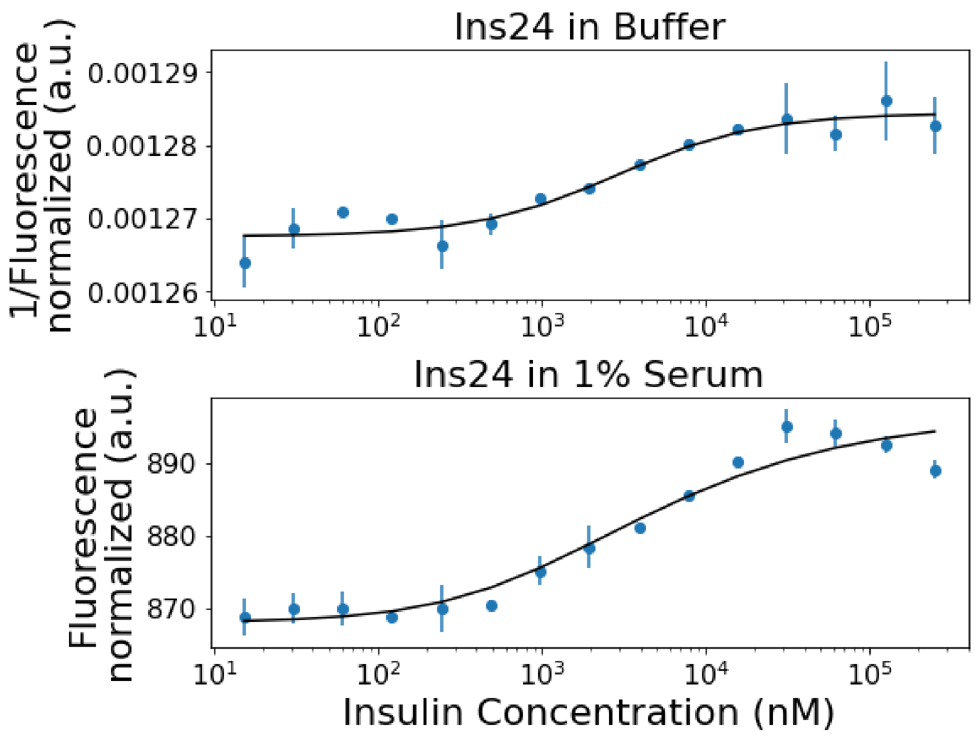


Microscale thermophoresis was used to measure insulin binding of ins24 in buffer (top) and 1% serum (right). F-modified, Cy5-labeled Ins24 was titrated with unlabeled insulin. Each measurement was performed in duplicate. We measured $K_D$ in buffer = 2.4 ± 0.4 µM and $K_D$ in 1% serum = 2.6 ± 0.6 µM. Data were fit to the quadratic binding equation[3]:

$$\frac{C_{AT}}{C_A} = Fraction\ bound = \frac{(C_T + C_A + K_D - \sqrt{(C_T + C_A + K_D)^2 - 4C_T C_A}}{2C_A}$$

where $C_A$= the constant concentration (5 nM) of Ins24, and $C_T$ = the concentration of unlabeled insulin. Note, the MST signal amplitude sign changes in serum compared to in buffer due to the changes in pH and ionic strength, so the y-axis is flipped in the buffer data for ease of comparison.[4]

#### III. Supplementary Tables

Table S1: DNA sequences used in this study

| Sequences for N2A2 | Sequence 5'-3' | Ref |
|---|---|---|
| Forward primer sequencing adaptor *,** | *TCG TCG GCA GCG TCA GAT GTG TAT AAG AGA CAG* HNN NNN NNN N AG CAG CAC AGA GGT CAG ATG | |
| Reverse primer sequencing adaptor | *GTC TCG TGG GCT CGG AGA TGT GTA TAA GAG ACA G* **GA ATT C** TT CAC GGT AGC ACG CAT AGG | |
| Fiducial mark (displayed on flow-cell)† | ACC GAC GGA ACG CCA AAG AAA CGC AAG G | |
| Fiducial mark complement (incubated during experiment) | /cy5/CCT TGC GTT TCT TTG GCG TTC CGT CGG T | |
| VEGF selection | | |
| V FP forward primer | CCT CTC TAT GGA CAC ACT ACC CT | |
| V RP reverse primer | CTG CAC TGC GTT CCT GAT ACC CT | |
| VEGF library | CCTCTCTATGGACACAC-TACCCT $(N)_{30}$ AGGGTA-TCAGGAACGCAGTGCAG | |
| Forward primer sequencing adaptor *,** | *TCG TCG GCA GCG TCA GAT GTG TAT AAG AGA CAG* NNNN CCT CTC TAT GGA CAC AC T ACC CT | |
| Reverse primer sequencing adaptor | *GTC TCG TGG GCT CGG AGA TGT GTA TAA GAG ACA G* **GA ATT C** CTG CAC TGC GTT CCT GAT ACC CT | |
| VEGF aptamer SL-2B (displayed on flow-cell) | AGC AGC ACA GAG GTC AGA TGA ATT GGG CCC GTC CGT ATG GTG GGT CCT ATG CGT GCT ACC GTG AA | [5] |
| Insulin selection | | |
| I FP forward primer | GCG CAT ACC AGC TTA TTC AAT T | |
| I RP reverse primer | GCC GAG ATT GCA CTT ACT ATC T | |
| Insulin library | GCG CAT ACC AGC TTA TTC AAT T – $(N)_{40}$ - AGA TAG TAA GTG CAA TCT CGG C | |
| Forward primer sequencing adaptor *,** | *TCG TCG GCA GCG TCA GAT GTG TAT AAG AGA CAG* HNN NGC GCA TAC CAG CTT ATT CAA TT | |
| Reverse primer sequencing adaptor | *GTC TCG TGG GCT CGG AGA TGT GTA TAA GAG ACA G* **GA ATT C** GC CGA GAT TGC ACT TAC TAT CT | |
| Sequences for flow cytometry analysis | | |

| | | |
|---|---|---|
| VEGF positive control (SL2B bt) | /5Biosg/CAA TTG GGC CCG TCC GTA TGG TGG GT | [5] |
| Insulin positive control (IGA3 bt) | /5Biosg/GGT GGT GGG GGG GGT TGG TAG GGT GTC TTC | [6] |

*H: Mixed base code for A,T,C only; **N: Mixed base code for A,T,C,G; †Fiducial mark sequence was randomly generated, with the restriction of not containing any Ts; *Underlined and italic* sequences show Illumina-defined sequencing primers; **Bold**: EcoRI cut site. A short random region was inserted between the Illumina sequencing primer and our aptamer forward primer sequence to ensure during sequencing that there is enough base diversity in the first 5 cycles of sequencing by synthesis for the instrument to deconvolute the different clusters of oligonucleotides.

Table S2: VEGF aptamer sequences assessed with N2A2

Three N2A2 experiments were performed with VEGF titrations: natural DNA, W-modified and Y-modified. For each experiment, we have provided the random region sequence of each aptamer and the number of times the sequence appeared on the flow-cell (copy number). We also provide the average fluorescence (a.u.) and standard deviation at each concentration of VEGF tested for each aptamer.

Table located at: https://github.com/sohlab/non-natural-aptamer-array

Table S3: W-modified VEGF aptamer candidates tested

| Sequence Name | Sequence 5'-3' (Random region only) | $K_d$ (nM) |
|---|---|---|
| VEGF-1 | CCTTTCCCGAACGTTGGCCGAACGGCATCG | 4.2 |
| VEGF-2 | TGAATCATCCGCTGGCCGAATGGCAGCGGG | 8.4 |
| VEGF-3 | GACAGGCATCCGAATGGCCGCCTGATATTA | 3.7 |
| VEGF-4 | GGCAAACGTCCGAATGGTTTTATATTAGGC | 3.3 |
| VEGF-5 | AAAATTTAGGTGCTTCTTTTTGTAAATATT | 8.1 |
| VEGF-6 | CCGAATGGGGTTTGAAGTTATGTGGATTGT | 5.1 |
| VEGF-7 | TGGTTTTACTTGTTCTAGTGTTGTAAACAA | 4.2 |
| VEGF-8 | CCCAGTAGGGTGGCAGTCAGGGAGTACATA | 5.5 |
| Full length VEGF-4 sequence tested by BLI | /5BiotinTEG/CCT CTC TAT GGA CAC ACT ACC CTG GCA AAC G/i5OctdU/CCGAA/i5OctdU/GG/i5OctdU/ /i5OctdU//i5OctdU//i5OctdU/A/i5OctdU/A/i5OctdU//i5OctdU/AGGCAGGG/i5OctdU/A/i5OctdU/CAGGAACGCAG /i5OctdU/GCAG | |

Random regions from selected VEGF aptamer candidates. Flow cytometry-based binding assays were performed for each W-modified aptamer (**Fig S6**). Curve-fitting was performed using nonlinear regression in Prism.

Table S4: Insulin aptamer sequences assessed with N2A2

One N2A2 experiment was performed with F-modified aptamers with insulin titrations in buffer and 1% serum. We have provided the random sequence of each aptamer and the number of times the sequence appeared on the flow-cell (copy number). We also provide the average fluorescence (a.u.) and standard deviation at each concentration of insulin tested for each aptamer.

Table located at: https://github.com/sohlab/non-natural-aptamer-array

Table S5: F-modified insulin aptamer candidates tested

Insulin aptamer candidates sequences identified from analysis of N2A2 experiment (SI Methods 1.11). Aptamers were chosen by high binding performance on the flowcell (Table S4), or by high family performance (Fig S10). This list includes sequences for which full binding curves were completed (Fig S11); Single point binding assays for additional aptamers with expected medium/poor binding performance are not included.

| Sequence Name | Sequence 5'-3' (Random region only) | Family |
|---|---|---|
| ins1 | TATGGTAGGTCAACAGCACGCGTGCCTCTCAACTGGATCTCA | N/A |
| ins2 | TCACAGACCATGTTTCGGTAATCCTGGTATTCGGTATTTATA | 1633 |
| ins7 | TTGACCACTGGAGGCTAGGAGAGGTATAATGCAGAGTCTACA | 68 |
| ins9 | TCGGTCGGCATGGTATCCCCTCAACAGGTATCGTCACTCCAA | 17 |
| ins10 | TGGAACAGGCCGCAACTGGAACCGGCGCCATCGCCTATACCA | 30 |
| ins13 | TATTTCTCTATTTGTTGTATTTTGGCGTTGGCCTTTTTGTTA | N/A |
| ins14 | TCCGCTCTTTATTGTTTCTTGTTCTGGTCGTGTTCCCATCTA | N/A |
| ins15 | TCTATTGCTCTTTTTCTTTCTATGTTATTCCTATATCAGCCA | N/A |
| ins23 | TTCAGCACGGAGACGAAGCGGCATTAGGGTAAGAGCGCCAAA | N/A |
| ins24 | TCGTCGAGAACTCCTCCGGCTCTAACTCCGATTATAGCCAAA | 6 |

**IV. Supplementary Code**

Custom Chemistry Modifications

Within the Miseq control software (C:/Illumina/ControlSoftware/Recipes/V3), recipes can be copied and modified. We made a copy of the v3_Miseq_Amplicon recipe folder, then modified the Chemistry.xml by adding or changing the following three definitions: CompleteCycleSel, PrimeAlt and Paired End Turnaround. We also modified the Reads.xml to use the new chemistry definitions.

CompleteCycleSel

```
<ChemistryDefinition Name="CompleteCycleSel">
      <ChemistryStep Description="Deblock" Repeat="1">
        <Temp Duration="0" Temperature="22" />
        <PumpToFlowcell ReagentName="PR2" AspirationRate="2000"
Solution="0" DispenseRate="7500" Volume="250" />
               <!-- NaOH Wash -->
        <PumpToFlowcell ReagentName="Alt2" AspirationRate="500"
Solution="0" DispenseRate="7500" Volume="500" />
        <DispenseAndWait DispenseRate="7500" Duration="300000" /
>
        <PumpToFlowcell ReagentName="PR2" AspirationRate="2000"
Solution="0" DispenseRate="7500" Volume="250" />
               <!-- NaOH Wash -->
        <PumpToFlowcell ReagentName="Alt2" AspirationRate="500"
Solution="0" DispenseRate="7500" Volume="250" />
        <DispenseAndWait DispenseRate="7500" Duration="300000" /
>
        <PumpToFlowcell ReagentName="PR2" AspirationRate="2000"
Solution="0" DispenseRate="7500" Volume="250" />
               <!-- Hyb buffer wash -->
        <PumpToFlowcell ReagentName="Alt2" AspirationRate="500"
Solution="0" DispenseRate="7500" Volume="500" />
        <DispenseAndWait Duration="300000" />
      </ChemistryStep>
      <ChemistryStep Description="Incorporation" Repeat="1" IsIn
corporation="true">
        <Temp Duration="0" Temperature="22" />
                <!-- Ligand incubation -->
        <PumpToFlowcell ReagentName="Alt2" AspirationRate="500"
Solution="0" DispenseRate="7500" Volume="500" />
        <DispenseAndWait Duration="300000" />
        <PumpToFlowcell ReagentName="Alt2" AspirationRate="250"
Solution="0" DispenseRate="7500" Volume="15" />
        <DispenseAndWait Duration="300000" />
        <PumpToFlowcell ReagentName="Alt2" AspirationRate="250"
Solution="0" DispenseRate="7500" Volume="15" />
```

```
        <DispenseAndWait Duration="300000" />
        <PumpToFlowcell ReagentName="Alt2" AspirationRate="250"
Solution="0" DispenseRate="7500" Volume="15" />
        <DispenseAndWait Duration="300000" />
        <PumpToFlowcell ReagentName="Alt2" AspirationRate="250"
Solution="0" DispenseRate="7500" Volume="15" />
        <DispenseAndWait Duration="300000" />
        <PumpToFlowcell ReagentName="Alt2" AspirationRate="250"
Solution="0" DispenseRate="7500" Volume="15" />
        <DispenseAndWait Duration="300000" />
        <PumpToFlowcell ReagentName="Alt2" AspirationRate="250"
Solution="0" DispenseRate="7500" Volume="15" />
        <DispenseAndWait Duration="301000" />
        <PumpToFlowcell ReagentName="Alt2" AspirationRate="250"
Solution="0" DispenseRate="7500" Volume="15" />
        <DispenseAndWait Duration="302000" />
        <Temp Duration="0" Temperature="22" />
                <!-- Pre-imaging buffer wash -->
        <PumpToFlowcell ReagentName="Alt2" AspirationRate="1000"
 Solution="0" DispenseRate="7500" Volume="2000" />
        <DispenseAndWait Duration="5000" />
      </ChemistryStep>
    </ChemistryDefinition>
```

PrimeAlt

```
<ChemistryDefinition Name="PrimeAlt">
      <ChemistryStep Description="Buffer Wash" Repeat="1">
        <PumpToFlowcell ReagentName="PR2" AspirationRate="2000"
Solution="0" DispenseRate="7500" Volume="250" />
      </ChemistryStep>
      <ChemistryStep Description="NaOHWash" Repeat="1">
        <Temp Duration="0" Temperature="22" />
        <PumpToFlowcell ReagentName="PR2" AspirationRate="2000"
Solution="0" DispenseRate="7500" Volume="250" />
              <!-- NaOH wash -->
        <PumpToFlowcell ReagentName="Alt2" AspirationRate="1500"
 Solution="0" DispenseRate="7500" Volume="1500" />
        <DispenseAndWait DispenseRate="7500" Duration="300000" /
>
        <PumpToFlowcell ReagentName="PR2" AspirationRate="2000"
Solution="0" DispenseRate="7500" Volume="500" />
      </ChemistryStep>
      <ChemistryStep Description="ECOR1" Repeat="1">
        <Temp Duration="0" Temperature="37" />
               <!-- EcoR1 buffer and comp strand -->
        <PumpToFlowcell ReagentName="C1" AspirationRate="2000" S
olution="0" DispenseRate="7500" Volume="150" />
```

```
        <DispenseAndWait Duration="600000" />
               <!-- EcoR1 HF -->
        <PumpToFlowcell ReagentName="Alt1" AspirationRate="2000"
 Solution="0" DispenseRate="7500" Volume="100" />
        <DispenseAndWait Duration="300000" />
               <!-- EcoR1 buffer and comp strand -->
        <PumpToFlowcell ReagentName="C1" AspirationRate="2000" S
olution="0" DispenseRate="7500" Volume="100" />
        <DispenseAndWait Duration="600000" />
               <!-- EcoR1 HF -->
        <PumpToFlowcell ReagentName="Alt1" AspirationRate="2000"
 Solution="0" DispenseRate="7500" Volume="100" />
        <DispenseAndWait Duration="300000" />
        <Temp Duration="0" Temperature="22" />
               <!-- NaOH wash -->
        <PumpToFlowcell ReagentName="Alt2" AspirationRate="500"
Solution="0" DispenseRate="7500" Volume="500" />
        <DispenseAndWait Duration="300000" />
      </ChemistryStep>
      <ChemistryStep Description="AtailRev" Repeat="1">
        <Temp Duration="0" Temperature="60" />
        <PumpToFlowcell ReagentName="PR2" AspirationRate="2000"
Solution="0" DispenseRate="7500" Volume="500" />
        <PumpToFlowcell ReagentName="CMS" AspirationRate="2000"
Solution="0" DispenseRate="7500" Volume="75" />
        <PumpToFlowcell ReagentName="PR2" AspirationRate="2000"
Solution="0" DispenseRate="7500" Volume="20" />
        <DispenseAndWait DispenseRate="7500" Duration="10000" />
      </ChemistryStep>
       <ChemistryStep Description="WaterWash" Repeat="1">
        <Temp Duration="0" Temperature="22" />
        <PumpToFlowcell ReagentName="PR2" AspirationRate="2000"
Solution="0" DispenseRate="7500" Volume="250" />
               <!-- Water Wash -->
        <PumpToFlowcell ReagentName="Alt2" AspirationRate="1000"
 Solution="0" DispenseRate="7500" Volume="1000" />
        <DispenseAndWait DispenseRate="7500" Duration="300000" /
>
      </ChemistryStep>
      <ChemistryStep Description="BufferWash" Repeat="1" IsIncor
poration="true">
        <Temp Duration="0" Temperature="22" />
                <!-- Azide click handle -->
        <PumpToFlowcell ReagentName="Alt2" AspirationRate="500"
Solution="0" DispenseRate="7500" Volume="500" />
        <DispenseAndWait Duration="300000" />
```

```
        <PumpToFlowcell ReagentName="Alt2" AspirationRate="250"
Solution="0" DispenseRate="7500" Volume="15" />
        <DispenseAndWait Duration="300000" />
        <PumpToFlowcell ReagentName="Alt2" AspirationRate="250"
Solution="0" DispenseRate="7500" Volume="15" />
        <DispenseAndWait Duration="300000" />
        <PumpToFlowcell ReagentName="Alt2" AspirationRate="250"
Solution="0" DispenseRate="7500" Volume="15" />
        <DispenseAndWait Duration="300000" />
        <Temp Duration="0" Temperature="22" />
        <PumpToFlowcell ReagentName="PR2" AspirationRate="2000"
Solution="0" DispenseRate="7500" Volume="2000" />
        <DispenseAndWait Duration="5000" />
      </ChemistryStep>
      <ChemistryStep Description="Wash" Repeat="1">
        <Temp Duration="0" Temperature="22" />
        <PumpToFlowcell ReagentName="PR2" AspirationRate="2000"
Solution="0" DispenseRate="7500" Volume="250" />
                <!-- NaOH wash -->
        <PumpToFlowcell ReagentName="Alt2" AspirationRate="250"
Solution="0" DispenseRate="7500" Volume="500" />
        <DispenseAndWait DispenseRate="7500" Duration="180000" /
>
        <PumpToFlowcell ReagentName="PR2" AspirationRate="2000"
Solution="0" DispenseRate="7500" Volume="250" />
                <!-- NaOH wash -->
        <PumpToFlowcell ReagentName="Alt2" AspirationRate="500"
Solution="0" DispenseRate="7500" Volume="250" />
        <DispenseAndWait DispenseRate="7500" Duration="180000" /
>
        <PumpToFlowcell ReagentName="PR2" AspirationRate="2000"
Solution="0" DispenseRate="7500" Volume="250" />
                <!-- Hyb buffer wash -->
        <PumpToFlowcell ReagentName="Alt2" AspirationRate="500"
Solution="0" DispenseRate="7500" Volume="500" />
        <DispenseAndWait Duration="300000" />
      </ChemistryStep>
      <ChemistryStep Description="IncorpFM" Repeat="1" IsIncorpo
ration="true">
        <Temp Duration="0" Temperature="22" />
                <!-- Ligand incubation -->
        <PumpToFlowcell ReagentName="Alt2" AspirationRate="700"
Solution="0" DispenseRate="7500" Volume="700" />
        <DispenseAndWait Duration="300000" />
        <PumpToFlowcell ReagentName="Alt2" AspirationRate="250"
Solution="0" DispenseRate="7500" Volume="15" />
        <DispenseAndWait Duration="300000" />
```

```
        <PumpToFlowcell ReagentName="Alt2" AspirationRate="250"
Solution="0" DispenseRate="7500" Volume="15" />
        <DispenseAndWait Duration="300000" />
        <PumpToFlowcell ReagentName="Alt2" AspirationRate="250"
Solution="0" DispenseRate="7500" Volume="15" />
        <DispenseAndWait Duration="300000" />
        <PumpToFlowcell ReagentName="Alt2" AspirationRate="250"
Solution="0" DispenseRate="7500" Volume="15" />
        <DispenseAndWait Duration="300000" />
        <PumpToFlowcell ReagentName="Alt2" AspirationRate="250"
Solution="0" DispenseRate="7500" Volume="15" />
        <DispenseAndWait Duration="300000" />
        <PumpToFlowcell ReagentName="Alt2" AspirationRate="250"
Solution="0" DispenseRate="7500" Volume="15" />
        <DispenseAndWait Duration="301000" />
        <PumpToFlowcell ReagentName="Alt2" AspirationRate="250"
Solution="0" DispenseRate="7500" Volume="15" />
        <DispenseAndWait Duration="302000" />
        <Temp Duration="0" Temperature="22" />
                 <!-- Pre-imaging buffer wash -->
        <PumpToFlowcell ReagentName="Alt2" AspirationRate="1000"
 Solution="0" DispenseRate="7500" Volume="2000" />
        <DispenseAndWait Duration="5000" />
      </ChemistryStep>
    </ChemistryDefinition>
```

<u>Paired End Turnaround</u>

```
<ChemistryDefinition Name="PairedEndTurnaround">
   <!-- Modified for KODXL Polymerase -->
 <ChemistryStep Description="Pre-resynthesis_tempramp">
      <PumpToFlowcell ReagentName="C2" AspirationRate="2000" Sol
ution="0" DispenseRate="7500" Volume="300" />
      <PumpToFlowcell ReagentName="PR2" AspirationRate="2000" Di
spenseRate="7500" Volume="120" />
      <Temp Temperature="55" />
    </ChemistryStep>
    <ChemistryStep Description="Resynthesis" Repeat="12" >
       <PumpToFlowcell ReagentName="LDR" AspirationRate="250" So
lution="0" DispenseRate="7500" Volume="75" />
        <Temp Temperature="75" />
        <Wait Duration="45000" />
               <!-- KodXL buffer -->
        <PumpToFlowcell ReagentName="C2" AspirationRate="250" So
lution="0" DispenseRate="7500" Volume="150" />
        <Temp Temperature="50" />
        <Wait Duration="30000" />
```

```
              <!-- KodXL Poly and dNTPs -->
        <PumpToFlowcell ReagentName="C3" AspirationRate="250" So
lution="0" DispenseRate="7500" Volume="75" />
        <Temp Temperature="75" />
        <Wait Duration="60000" />
    </ChemistryStep>
    <ChemistryStep Description="Post-resynthesis_wash">
      <PumpToFlowcell ReagentName="PR2" AspirationRate="2000" Di
spenseRate="7500" Volume="120" />
      <Temp Temperature="20" Duration="45000" />
    </ChemistryStep>
    <ChemistryStep Description="Linearisation 2">
       <PumpToFlowcell ReagentName="LMX2" AspirationRate="2000"
Solution="0" DispenseRate="7500" Volume="300" />
        <PumpToFlowcell ReagentName="LMX2" AspirationRate="250"
Solution="0" DispenseRate="7500" Volume="75" />
      <Temp Temperature="38" />
        <Wait Duration="300000"/>
        <PumpToFlowcell ReagentName="LMX2" AspirationRate="250"
DispenseRate="7500" Volume="25" />
  <Wait Duration="300000" />
        <PumpToFlowcell ReagentName="LMX2" AspirationRate="250"
DispenseRate="7500" Volume="25" />
        <Wait Duration="300000" />
        <PumpToFlowcell ReagentName="PR2" AspirationRate="2000"
Solution="0" DispenseRate="7500" Volume="120" />
    </ChemistryStep>
  </ChemistryDefinition>
```


Reads.xml


```
<?xml version="1.0"?>
<ReadDefinitions Version="Fraise 0.7.2 Amplicon v1.1">
    <ReadDefinition Name="FirstRead">
      <FirstBase ChemistryName="FirstBase" Cycles="1" />
      <Imaging ChemistryName="CompleteCycle" Cycles="NumCycles -
 1" />
      <End ChemistryName="EndRead" />
    </ReadDefinition>
    <ReadDefinition Name="IndexRead1" IsIndexed="true" >
      <FirstBase ChemistryName="FirstBase" Cycles="1" />
      <Imaging ChemistryName="CompleteCycle" Cycles="NumCycles -
 1" />
      <End ChemistryName="EndRead" />
    </ReadDefinition>
    <ReadDefinition Name="SecondRead" IsIndexed="false">
      <FirstBase ChemistryName="FirstBase" Cycles="1" />
```

```
      <Imaging ChemistryName="PrimeAlt" Cycles="1" />
      <Imaging ChemistryName="CompleteCycleSel" Cycles="NumCycle
s - 2" />
      <End ChemistryName="EndRead" />
    </ReadDefinition>
    <ReadDefinition Name="IndexRead2" IsIndexed="true">
      <Imaging ChemistryName="CompleteCycle" Cycles="NumCycles"
/>
      <End ChemistryName="EndRead" />
    </ReadDefinition>
  </ReadDefinitions>
```
